# Cross-linking mass spectrometry discovers, evaluates, and validates the experimental and predicted structural proteome

**DOI:** 10.1101/2022.11.16.516813

**Authors:** Tara K. Bartolec, Xabier Vázquez-Campos, Alexander Norman, Clement Luong, Richard J. Payne, Marc R. Wilkins, Joel P. Mackay, Jason K. K. Low

## Abstract

Significant recent advances in structural biology, particularly in the field of cryo-electron microscopy, have dramatically expanded our ability to create structural models of proteins and protein complexes. However, many proteins remain refractory to these approaches because of their low abundance, low stability or – in the case of complexes – simply not having yet been analysed. Here, we demonstrate the power of combining cross-linking mass spectrometry (XL-MS) with artificial intelligence-based structure prediction to discover and experimentally substantiate models for protein and protein complex structures at proteome scale. We present the deepest XL-MS dataset to date, describing 28,910 unique residue pairs captured across 4,084 unique human proteins and 2,110 unique protein-protein interactions. We show that integrative models of complexes driven by AlphaFold Multimer and inspired and corroborated by the XL-MS data offer new opportunities to deeply mine the structural proteome and interactome and reveal new mechanisms underlying protein structure and function.

## INTRODUCTION

Proteins are the primary effectors in biology. Their function is determined in large part by their three-dimensional structure and by the protein-protein interactions (PPIs) that they form. A system-wide understanding of protein structure and interactions has thus been a long-standing goal.

To this end, the protein-protein interactome has been systemically catalogued using several approaches, including yeast two-hybrid (Y2H), affinity-purification mass spectrometry (AP-MS) and by the Bio-ID proximity assay and variants (reviewed in (1)). These methods have been very successful in identifying both direct and indirect protein interactions and at least one interactor has been curated for almost 50% of the human proteome (2). However, these data do not provide structural or mechanistic information on interactions and do not necessarily analyse proteins interacting in their native state. These significant shortcomings call for additional strategies to better define the interactome.

Despite intensive efforts in the fields of X-ray crystallography, nuclear magnetic resonance (NMR) and cryo-electron microscopy (cryo-EM), only 35% of human proteins have any representation in the Protein Data Bank (PDB) (3, 4). Many are only partially resolved: only 17% of residues in human proteins are present in the PDB and experimental structures exist for only 6% of known human protein-protein interactions (5). Furthermore, the heterologous overexpression and purification of proteins are often necessary to produce enough material for these techniques, with only 5% of the human proteins in the PDB produced from native sources (3, 4). This situation raises possible concerns about the integrity of structures and complexes that are generated using such approaches.

*In lieu* of experimental structures, machine learning-based structural modellers such as AlphaFold (6) have been shown to be highly accurate under controlled tests (7) and have greatly expanded the coverage of structural proteomes across a range of species (8, 9). However, because the training dataset (the PDB) has a very low representation of truly native structures, the accuracy of these models and the question of how to assess them remain to be determined.

Cross-linking mass spectrometry (XL-MS) provides a means to assay native protein structure and protein-protein interactions in a parallel fashion (reviewed in (10, 11)). Chemical cross-linkers containing at least two reactive groups are introduced into a protein sample to covalently link amino acids within spatially constrained reactions. The identification of cross-linked peptides via mass spectrometry enables the definition of the linked residues, providing conformational constraints for a protein chain or PPI interface. These constraints, despite having low effective resolution, can be used to validate experimental structures, help inform modelling of proteins with unknown structures, and guide docking studies of PPIs (reviewed in (10)). Taking into consideration the relatively low protein sample requirements, XL-MS can provide a first view of the structures and interactions of many poorly studied or less accessible proteins (12).

Recent technical developments in XL-MS have established its use at scale and in native contexts such as in intact human cells (13, 14) and even mammalian tissues (15). However, the complexity and dynamic range of the proteome and interactome leads to under-sampling. One means to achieve better proteome coverage whilst preserving *in vivo* or near-*in vivo* proteoform states is to isolate and then cross-link intact organelles (16, 17).

Furthermore, the use of diverse cross-linker reactivities can increase the density of structural constraints by sampling a larger protein sequence space (18, 19).

To establish the utility of large-scale XL-MS for the characterisation of the structural proteome and interactome, we have generated a high-coverage and high-density cross-link dataset for a cultured human cell line. Fractionated organelles were cross-linked with three different chemistries (DHSO (20), DSSO (21), DMTMM (22)), and a multi-step analytical pipeline used to extensively fractionate and identify cross-linked peptides. We identified 91,709 cross-link spectral matches (CSMs) representing 28,910 unique residue pairs across 4,084 proteins and 2,110 protein-protein interactions. This resource is the largest reported to date for any species.

Benchmarking against the protein structure and PPI literature demonstrates the integrity and utility of our dataset, which provides the first orthogonal validation for many high resolution (but *in vitro*) experimental structures. In parallel, our cross-links also identify new PPIs and confirm many PPIs that were previously only identified using systems-level approaches. Importantly, our data also provide new structural information for a broad range of proteins and PPIs, including proteins that lack any experimental characterisation. We demonstrate how cross-links can be utilised to evaluate, validate, and support structure modelling platforms such as AlphaFold. Overall, we conclude that XL-MS in combination with structural modelling pipelines can produce highly accurate models to extend our understanding of the structural proteome, including in the context of the interactome.

## RESULTS

### Orthogonal cross-linkers, sample enrichment and improved database search strategies generate the deepest cross-linked proteome to date

To provide deep coverage of the human structural proteome, we performed crude subcellular fractionation of HEK293 cells and cross-linked these fractions in two reactions with three orthogonal cross-linkers: DHSO (20), DSSO (21), DMTMM (22). Following protein digestion, we fractionated the resulting peptides using sequential offline size exclusion chromatography and high pH reversed-phase liquid chromatography (**Figure 1A**).

**Figure 1.**
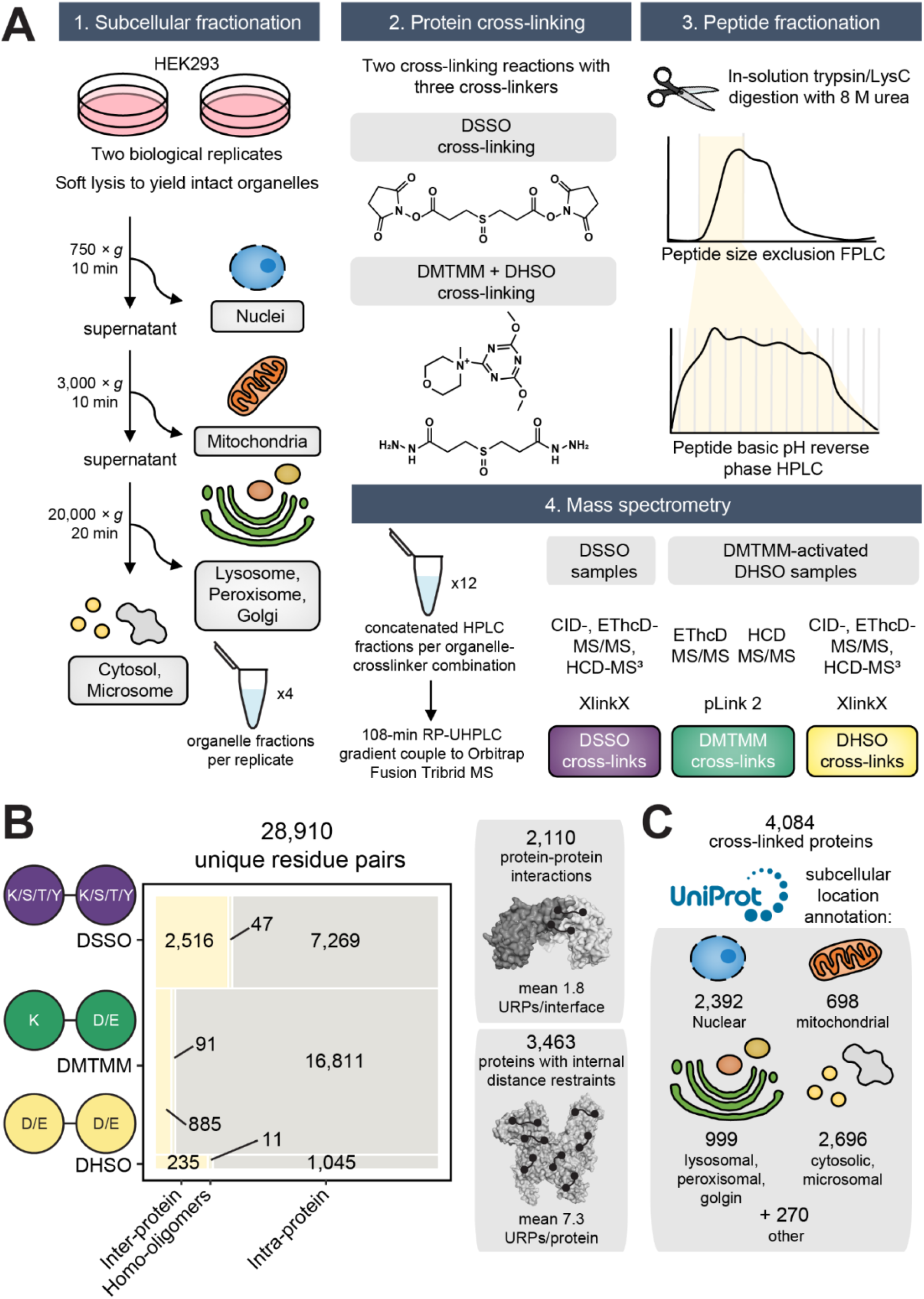
Generation of a deep human cross-linking mass spectrometry dataset. **(A)** Experimental overview. **(B)** Breakdown and summary statistics of the cross-linked unique residue pairs (URP) and the underlying captured unique protein-protein interactions or internally-linked proteins. Inter-protein refers to cross-linking between peptides mapping to two distinct UniProt accessions, intra-protein to the same accession, and homo-oligomer to same accession but with overlapping peptide sequences (and hence must derive from two separate molecules). **(C)** The size and breakdown of annotated subcellular localisations detected in the cross-linked proteome. Note that redundancies are caused by proteins with multiple annotated subcellular assignments.

From mass spectrometry, we identified 4,084 unique proteins linked by 28,910 unique cross-linked residue pairs (which we define as ‘unique residue pairs’, or URPs) from 91,709 crosslink spectral matches (CSMs) (**Figure 1B, C; Supp Table 1**). These URPs can be further classified into 3,785 and 25,125 inter-protein (including homo-oligomers) and intra-protein URPs, respectively.

Our URP list was generated using stringent quality control measures (**Methods**, **Supp Figure 1,** (16)). These enabled the control of the false discovery rate (FDR) to ≤1% at the URP level for both intra-protein and inter-protein links, and 1.9% at the protein-protein interaction (PPI) level for inter-protein links. All three cross-linkers produced inter-to intra-protein URP ratios in line with theoretical expectations for their maximum distance constraints (23), indicating an appropriately controlled FDR.

To our knowledge this is the largest cross-linking mass spectrometry (XL-MS) dataset to date, containing more than twice the number of URPs of the next largest studies (14, 24, 25). Protein abundance data from the PaxDB database (26) revealed that we sampled the proteome at a significantly deeper level than previous work. We identified 1.6-fold more proteins than the previous benchmark (14), and the median abundance of a cross-linked protein in our dataset was 1.8-fold lower (**Supp Figure 2A**). Furthermore, the density of cross-links defining each protein-protein interaction and intra-linked protein was high **(Supp Figure 2B)**. Overall, these demonstrate that the dataset is deep and of high quality.

Most proteome-wide XLMS studies to date have relied on N-hydroxysuccinimide-based (NHS) cross-linkers, which primarily target the ε-amino side chain of lysine (K) residues (21). Although reactions with the hydroxyl side chains of serine (S), threonine (T) and tyrosine (Y) are also possible (27, 28), they are often not considered due to the significant (and in most cases, prohibitive) increase in computation time during peptide database searches. In contrast to most other comparable studies, we included possible cross-links to S/T/Y during in our analysis of DSSO spectra. This strategy yielded 1.3-fold more DSSO URPs compared to K-K cross-link searching alone (9,832 vs 7,829 URPs). Linkages involving S/T/Y residues made up 24% of all DSSO URPs (**Supp Figure 3A**), consistent with previous smaller scale studies (27, 28). Many S/T/Y linked residues were initially localised to a nearby K residue within the same peptide **(Supp Figure 3B)**. For many of these peptides, their identification scores significantly improved when S/T/Y reactivity were considered (**Supp Figure 3C**), indicating an improved accuracy in the localisation of cross-linking sites. It also enabled us to identify ~20% more PPIs than a K-K-only search strategy (1,464 rather than 1,243 PPIs).

Our data indicate that different cross-linker chemistries might be better suited to certain subcellular niches. The proportion of URPs arising from each cross-linker varied significantly between organellar fractions (nucleus, cytoplasm, mitochondria and Golgi). DSSO was the most effective at cross-linking the nuclear fraction, whereas DHSO was most effective for the mitochondrial and Golgi fractions (**Supp Figure 4A**). These differences might reflect variation in amino acid composition, local pH or even the solubility or permeability of a cross-linker in a given compartment. We also noted that the nuclear fraction produced a significantly higher proportion of inter-protein URPs than any other fraction, suggesting denser protein packing or better preservation of protein complexes (**Supp Figure 4B**).

### Benchmarking against experimental structures demonstrates the high quality of the dataset

We next used structures from the PDB (4) to assess the quality of our data. Forty-three percent (9,152) of our 21,367 unambiguous URPs (*i.e*., URPs from cross-linked peptides for which each peptide sequence could be uniquely mapped to a single UniProt accession code) could be mapped onto 10,332 unique experimental structures (**Supplementary Table 2**). Euclidean distances (Cα-Cα) were calculated using Xwalk (29) and URPs considered satisfied if their minimum mapped distance on any PDB entry was within the maximum theoretical distance of 30 Å for DHSO (20) and DSSO (21), and within 25 Å for DMTMM (22, 30).

Considering only intra-chain URPs, 7,860 URPs mapped onto 10,110 structures of 1,406 proteins, of which 99% (DHSO), 97% (DSSO), and 89% (DMTMM) were satisfied (**Figure 2A**, **Supplementary Table 2**). In contrast, randomly sampled residue pairs within each structure met the distance cut-offs in only ~40–60% of cases (**Figure 2A**). For inter-chain linkages (including homo-dimeric URPs from overlapped peptide sequences), 1,292 URPs describing 519 unique PPIs were mapped onto 2,274 distinct PDB entries. We observed slightly lower distance satisfaction rates of 90% for DSSO, 72% for DMTMM, and 96% for DHSO (consistent with previous work (31)), whereas randomly sampled inter-chain URPs had very low satisfaction rates (~7–12%).

**Figure 2.**
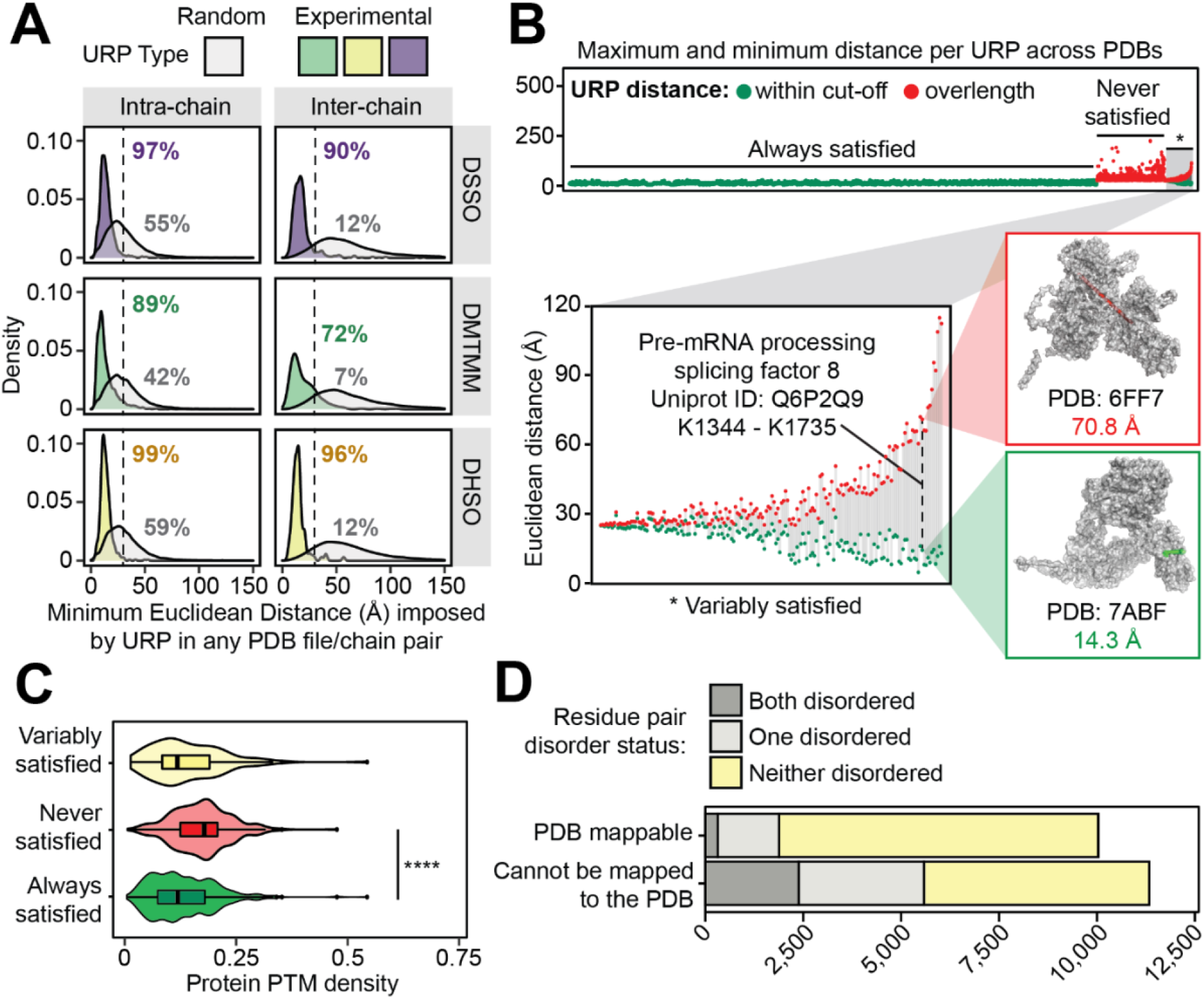
Cross-link distance constraints are validated by, and uniquely contextualise the variability within, *in vitro* experimental structures curated in the Protein Data Bank. **(A)** The distribution of minimum Euclidean distances (Å, Cα-Cα) imposed by unique residue pairs (URPs) across redundant possible chain-pairs and/or Protein Data Bank (PDB) entries. Only ‘unambiguous’ URPs are considered in these analyses, which are those with underlying cross-linked peptide sequences that were uniquely mapped to a single protein sequence in the canonical human proteome. Random residue pairs with appropriate sidechain reactivities were simulated for each individual PDB structure. The percentage of URPs falling within the cross-linker specific distance cut-offs (dotted lines, 25 Å for DMTMM or 30 Å for DHSO/DSSO) are also indicated. **(B)** The range of Euclidean distances imposed by URPs mapped across multiple unique PDB entries, considering only PDB entries with one possible chain-pair configuration for the URP. The minimum and maximum distances observed for each URP are plotted in-line as circles joined by a grey line. Green fill denotes a structure for which the distance in question is falls within the relevant cut-off, and red fill denotes a structure in which the distance violates the cut-off. The *top* panel shows the range distribution for all unambiguous URPs, stratified by their global satisfaction rate and ordered by increasing difference in distances within these categories. The distribution of the URP subset with variable satisfaction across PDB entries (*), is shown on the *lower left*. Inset structures to the right show an example of a variably satisfied URP from the pre-mRNA-processing-splicing factor 8 protein (Q6P2Q9), indicated by the dashed line. **(C)** The density of PhosphoSitePlus-annotated post-translational modifications (number of distinct annotated modification sites/length of protein) of cross-linked protein(s) (with the maximum value for the two proteins used for inter-protein links) for each URP mapping stratification type (by variability of URP satisfaction as described above). **** is p = 2.2 x 10^-16^, from a one-tailed Wilcoxon rank sum tests with continuity corrections. **(D)** The distribution of unambiguous URPs involving residues from regions predicted to be disordered, stratified by whether the URPs are resolved in PDB structures.

It is also notable that our cross-linking was carried out under significantly more native-like conditions than those typically used for the determination of protein structures. Structures are generally determined from heterologously expressed polypeptides that often comprise only a fraction of the full protein sequence and are determined in the absence of interactions with partners. The remarkably high overall satisfaction rates we observe therefore provide large-scale experimental corroboration of thousands of *in vitro* experimental structures.

### Cross-links capture and confirm alternative structural conformers

We noted many URPs that could map to more than one PDB structure for the same protein and therefore asked whether our data could provide insights into possible conformational plasticity of these proteins. We examined 4,857 URPs that could be mapped onto multiple PDB structures and compared the minimum and maximum Euclidean distances for these URPs across all of their PDB entries (**Figure 2B**, **Supplementary Table 2**). Most URPs (85.4%, ‘Always satisfied’ in **Figure 2B top panel**) were always mapped within the cross-linker cut-off distances regardless of PDB structure.

In contrast, although 529 URPs were not satisfied in any available structure, we observed 178 URPs that were satisfied in some but not all structures (‘Never Satisfied’ or ‘Variably Satisfied’, respectively, **Figure 2B)**. For example, the URP K1344-K1735 in the pre-mRNA splicing factor 8 (Uniprot: Q6P2Q9) differed dramatically in distance depending on which specific precursor subcomplex of the spliceosome it was mapped onto. The two residues show a distance of 71 Å in the structure of a spliceosome core variant that contains the U2/U6 catalytic RNA network (PDB: 6FF7 (32)), whereas the same residues are 14 Å apart in a structure lacking these RNAs (PDB: 7ABF, (33), **Figure 2B inset**). In another example, the URP E673-K890 in the DNA replication licensing factor MCM2 (UniProt: P49736) shows a distance of 10 Å when the complex that it is a part of – the CDC45-MCM-GINS helicase – is not engaged in the replisome (PDB: 6XTX (34)) and 39 Å when it is engaged in the replisome (PDB: 7PFO (35)) (**Supp Figure 5**). These observations suggest that URPs that are not satisfied in currently available protein structures might in some cases flag the existence of alternative conformations or architectures that are yet to be experimentally characterised.

Alternative conformations can arise from the formation of distinct complexes, as above, or from changes in post-translational modifications (PTMs). We observe that the never satisfied URPs tended to fall in proteins that are more densely post-translationally modified according to the PhosphoSitePlus database (36) compared to those satisfied in all PDB structures (**Figure 2C**; p < 2.2 × 10^-16^). This finding underscores the fact we are probing endogenous proteins in a near-cellular environment, and therefore probably capturing protein conformations and complex architectures that more closely reflect the *in vivo* state than do experimental high-resolution structures of isolated polypeptides expressed from heterologous systems (*e.g. E. coli*) that do not possess relevant human PTMs.

As noted above, we were unable to map more than half (~53%) of our unambiguous URPs due to the absence of cognate experimental structures. Because intrinsically disordered regions (IDRs) are resistant to structure determination, we investigated whether our unmappable URPs resided in such regions. Surprisingly, more than half of the URPs without PDB resolution did not lie in a disordered region, as annotated by a consensus of predictions curated in MobiDB (37) (**Figure 2D**). These URPs therefore likely define experimental distance restraints for ordered regions within structures that are either difficult to resolve (because of conformational dynamics, recalcitrance to purification efforts or highly contextual conformations) or simply involve unstudied proteins.

### Cross-links enable the high-throughput experimental assessment of protein structure predictions

Next-generation structural modelling programs such as AlphaFold2 (6) have generated enormous recent interest because of their performance in controlled environments such as CASP14 (7). However, the accuracy of such structural predictions for proteins in their native cellular context is less well-established. The near-native structural constraints provided by our cross-link resource provides a unique opportunity to address this question and we therefore asked how well our unambiguous intra-protein URPs map onto the recent AlphaFold2 (AF2) predictions of the human proteome (9). We only considered URPs that mapped to ‘high-confidence’ residues – defined by a value of ≥ 70 for the AlphaFold pLDDT quality metric. Despite this restriction, the AF2 dataset significantly increased the number of unambiguous URPs that we could map, with 12,359 URPs able to be resolved across 2,467 unique proteins (**Figure 3A, Supplementary Table 3**). This represents a 1.4-fold increase in resolved URPs compared to our PDB mapping above, and an increase of 1.8-fold in the number of proteins to which URPs could be mapped.

**Figure 3.**
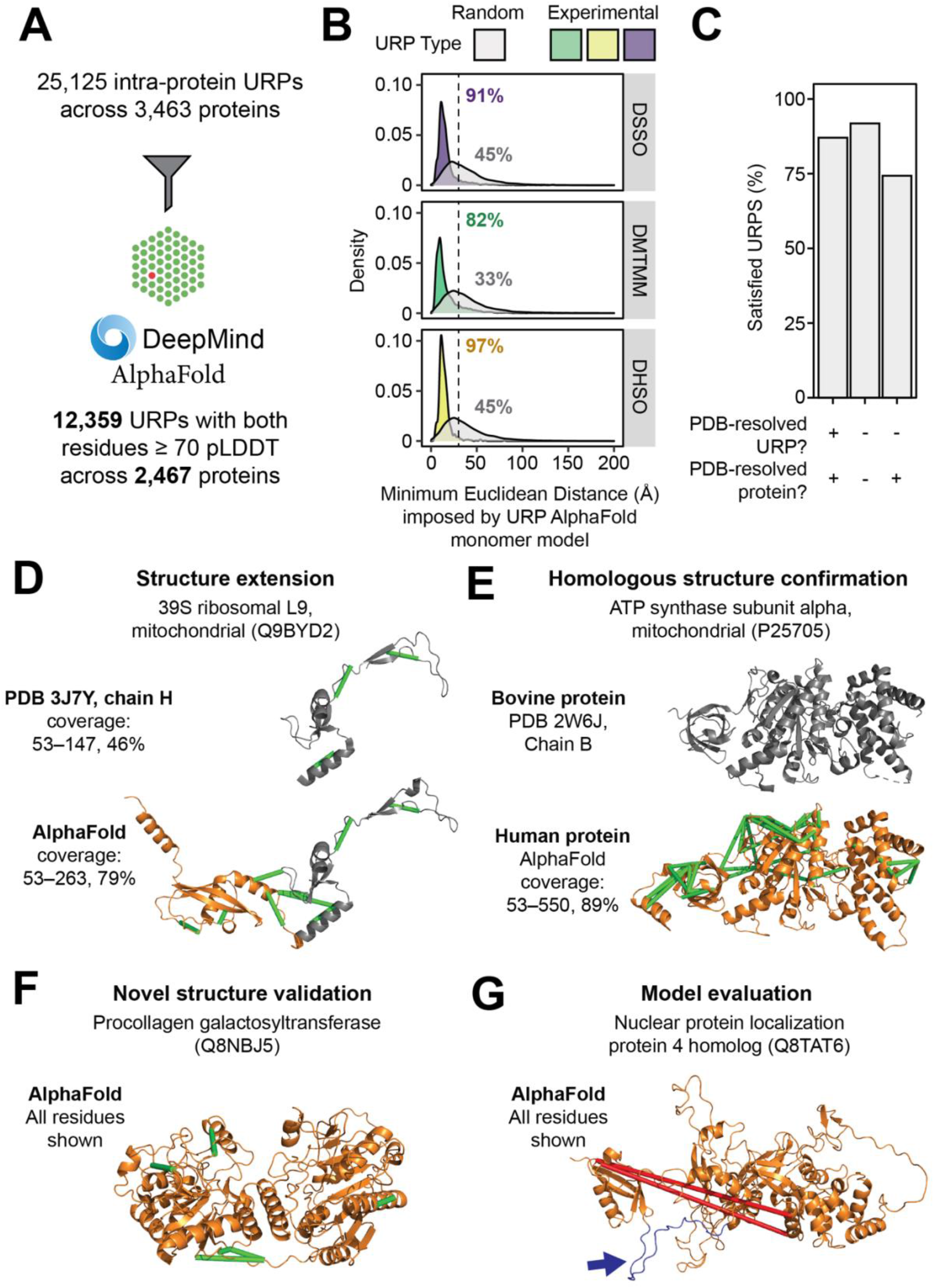
The high-throughput experimental assessment of AlphaFold monomeric protein structure predictions using cross-link distance constraints. **(A)** Statistics for the mapping of unique URPs onto AlphaFold 2 (AF2) monomer models. Only high-confdence URPs (both residues within pLDDT ≥ 70) were used for further analyses. **(B)** Fulfillment of URPs on AF2 models. For each AF2 monomer model, five random URPs were generated for each cross-linker with appropriate side-chain reactivities. The distribution of Euclidean distances (Å, Cα −Cα) determined for experimental and random URPs is shown, stratified by cross-linker. Annotations show the percentage of URPs fulfilled for each subset. The cut-off for each cross-linker is indicated by a dotted line (25 Å for DMTMM or 30 Å for DHSO/DSSO). **(C)** Overall URP satisfaction rate across URPs with different degrees of PDB resolution. **(D-G)** Examples of the use of URPs for model validation and evaluation. Protein names are official UniProt entry names. The range of residues shown are indicated to the left of each structure. Green URPs are satisfied, red URPs are overlength. **(D)** The model for Q9BYD2 shows an example of a PDB-resolved human protein that has benefitted from increased structural coverage (*orange*) and is now corroborated by nine experimental URPs. Grey regions within structures represent those derived from experimental PDB structures. **(E-F)** show proteins without any PDB entry for the human proteins (*orange*). Grey regions within structures represent those derived from experimental PDB structures. **(E)** The AF2 model of the mitochondrial ATP synthase subunit alpha with a known bovine structural homologue, 44 URPs support this model. **(F)** The AF2 model of the procollagen galactosyltransferase (Q8NBJ5) that does not have any PDB entries for itself or of homologous proteins. Five URPs support this model. **(G)** An example where the two identified cross-links do not fit the AF2 model. However, while the cross-linked residues are within well-modelled domains, the two domains are separated by a low-confidence, disordered loop (coloured in *blue* and highlighted with a blue arrow).

The distance distribution and satisfaction rates of these high-confidence URPs on the AF2 models (**Figure 3B**, **Supp Figure 6A**) were comparable to the values observed for intra-protein URPs mapped to PDB structures (**Figure 2A**), spanning 82–97% satisfaction across cross-linkers, compared to ~33-45% for randomly sampled URPs. This observation confirms that the predictions made by AF2 are of very high quality.

Given that AF2 is trained on the PDB, we next asked whether URP satisfaction rates on AF2 models varied depending on whether or not an experimental structure also existed for that protein (**Supplementary Table 3**). We first stratified the high-confidence URPs into three categories based on their PDB status at both the URP level and the overall protein level (**Figure 3C**). URPs for which both residues could be observed in a PDB structure had an 87% satisfaction rate on their corresponding AF2 model. Unexpectedly, URPs in proteins for which no experimental structure is currently available had even better satisfaction rates (92%). Furthermore, URPs from proteins for which a structure is available – but one that does not encompass the cross-linked residues – had a substantially lower satisfaction rate of 74%. These URPs might capture protein regions that display significant conformational dynamics, highlighting the fact that each AF2 model captures only a single snapshot of a protein’s conformational landscape.

Our data demonstrate the potential for integrated systems-wide approaches (such as the combination of AF2 and XL-MS) to address known biases of structural proteome coverage in the PDB – for example, towards well-behaved and highly ordered domains/proteins.

### Cross-links experimentally corroborate hundreds of AlphaFold models of proteins with unknown structures

As we mapped our URPs to the AF2 models, we noted that some models increased the structural coverage of a PDB-resolved protein. For example, the L9 subunit of the mitochondrial 39S ribosome (Uniprot: Q9BYD2) is partially resolved in several PDB entries; for example, in PDB: 3J7Y (38) residues 53–147 are observable, and we can map three URPs to this region, all of which are satisfied (**Figure 3D, *top***). However, the AF2 model resolved an additional 116 ordered C-terminal residues compared to the experimental structure, all of which have pLDDT scores of ≥ 70. An additional six URPs mapped to this region (including three that bridge the PDB-resolved and AF2-only regions) and all were satisfied in the AF2 model (**Figure 3D, *bottom***). This agreement provides strong corroboration for the predicted model and in this context, we reiterate that the cross-linking data were obtained in a native-like context, giving additional confidence that this is the relevant structure in the environment of an intact ribosome *in vivo*.

Of the set of 737 proteins that were absent from the PDB but for which cross-links were observed, 624 had all of their high-confidence URPs satisfied in the corresponding AF2 model (**Supp Figure 6B, Supplementary Table 3**). Our dataset provides experimental validation for these models in one of several ways. Firstly, there were 268 instances where the structure of the human protein is unknown, but a structure exists for a homologue. This included the alpha subunit of mitochondrial ATP synthase (Uniprot: P25705). The structure of the human protein has not been reported but the structure of the bovine protein (Uniprot: P19483) (97% sequence identity) is resolved in PDB: 2W6J (**Figure 3E**). Not surprisingly, the AF2 model for the human protein closely resembles the bovine structure, and our set of 44 experimental URPs verifies the conserved fold.

Secondly, there were 356 proteins for which we observed cross-links that were both absent from the PDB and also displayed a very low degree of structural precedent. For example, the procollagen galactosyltransferase 1 enzyme (Uniprot: Q8NBJ5) has neither an existing PDB entry nor a PDB structure for a homologous protein (using protein BLAST against the PDB database (39)). AF2, however, produced a model with over 85% of residues having pLDDT ≥ 70 (**Figure 3F**). Importantly, all five high-confidence URPs were satisfied within this model, compared to only 8% in the random URP control. Similarly, the very-long-chain 3-oxoacyl-CoA reductase (Uniprot: Q53GQ0) had little structural precedent but AF2 produced a model with >95% of residues having pLDDT ≥ 70 and for which all nine URPs were satisfied (**Supp Figure 6C**).

Thirdly, cross-links that do not corroborate AF2 models are also of interest as they indicate incongruence between experimental data and modelled structures. In total, there were 66 proteins with structure predictions for which all cross-links were violated in the AF2 model. For example, despite > 90% of the sequence the nuclear protein localisation protein 4 (NPL4; Uniprot: Q8TAT6) being well-modelled by AF2, neither of the two mappable URPs were satisfied on the model (**Figure 3G**). Examination of the model revealed that a ~20-residue linker with low pLDDT scores separates an N-terminal domain from the bulk of the protein and that both cross-links bridge these two domains. Repositioning of the N-terminal domain to satisfy the cross-links would involve a straightforward rigid-body rearrangement. Proteome-wide XL-MS data thus provide a resource to aid interpretation of – and potentially improve upon – structural models, making use of either manual ‘structural sculpting’ (*e.g*., as implemented in XMAS (40)) or integrative modelling pipelines (reviewed in (41)) that can be driven by cross-linking restraints.

### Cross-links reveal and corroborate thousands of protein-protein interactions

Our cross-link resource identifies and provides insights into the interfaces of thousands of protein-protein interactions (PPIs). A total of 3,785 inter-protein URPs describe 2,110 PPIs, including 84 unambiguous homo-oligomers (**Figure 4A**).

**Figure 4.**
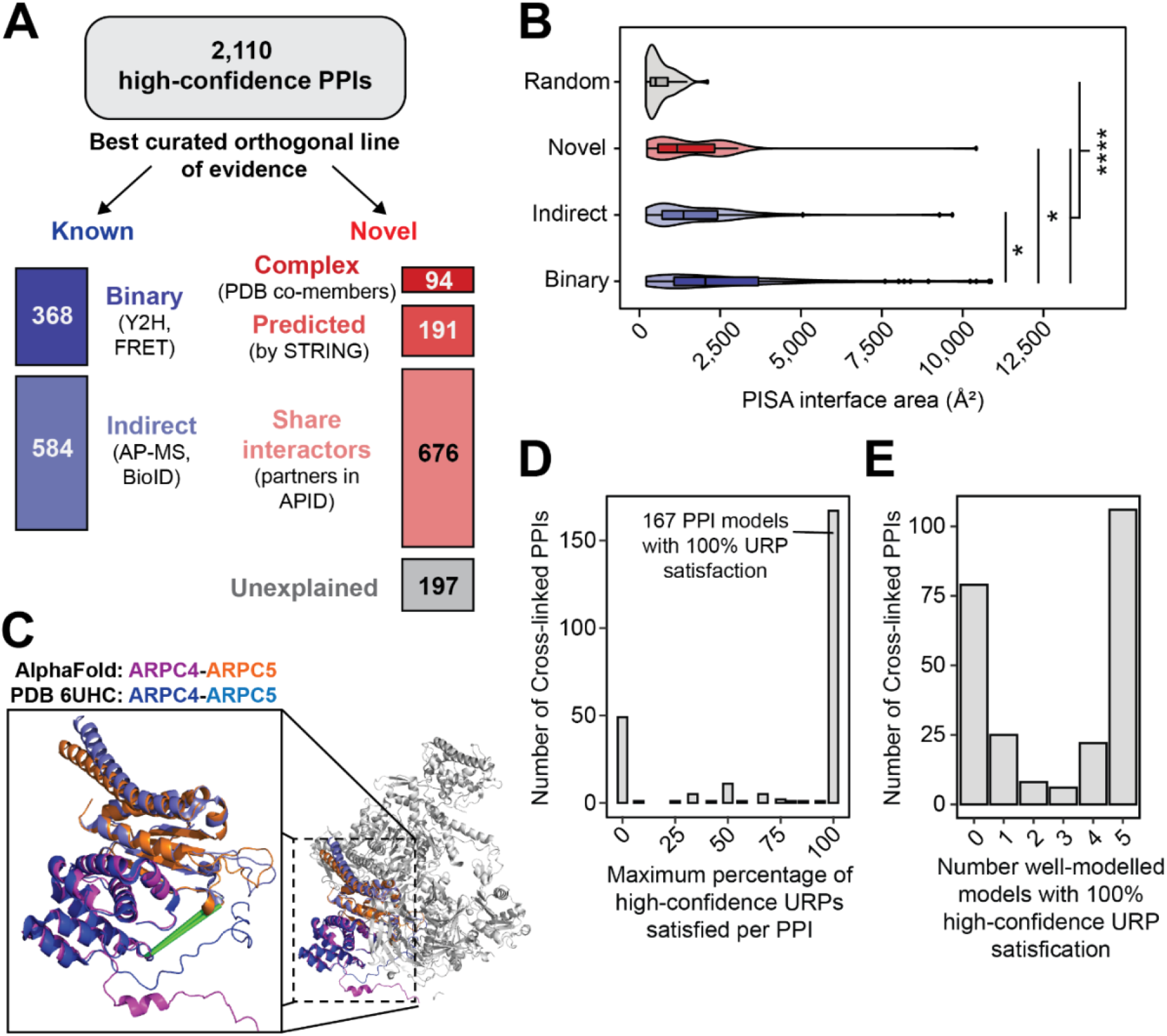
Discovery and AF Multimer-v2 modelling of protein-protein interactions. **(A)** Stratification of protein-protein interactions by their best existing curated evidence. Categories for ‘known’ PPIs include those annotated with a binary, or if not, an indirect interaction mapping technique in the APID database. Categories for ‘novel’ PPIs include proteins co-curated in the same CORUM complex, or PDB entry, or those predicted by STRING (combined scores at least 0.4), or those that share local interactome partners. **(B)** The largest interface surface area per PPI calculated by PISA for well-modelled AlphaFold Multimer-v2 models (clashscores ≤ 100, average interface residue pLDDT ≥ 70, interface areas > 200 Å^2^) generated for interactions with at least 2 URPs (stratified by best evidence – Binary, Indirect or Novel) and for random protein pairs. **(C)** The predicted AF Multimer-v2 model of the ARPC4-ARPC5 overlays very well (RMSD = 0.9 Å) with the known PDB: 6UHC structure of the full 7-subunit Arp2/3 complex. **(D and E)** Only high-confidence URPs (both cross-linked residues with pLDDT ≥ 70) were used for assessment. **(D)** The distribution of maximum percentage satisfaction rates of high-confidence URPs per well-modelled PPI. **(E)** The number of well-modelled AF Multimer-v2 models with fully-satisfied URPs per PPI.

Comparison of our dataset to the APID PPI meta-database (2) showed that 55% (1,158) of these PPIs had not been previously described (**Supplementary Table 4**). We therefore assessed the degree of orthogonal evidence available for each protein pair by sorting the PPIs according to their highest level of supporting evidence. Strikingly, 584 (more than 60%) of our 952 known interactions had only been described by indirect interaction mapping methods, such as affinity purification mass spectrometry and proximity ligation assays.

Of the novel interactions, most could be explained by leveraging existing systems-level annotations **(Figure 4A)**. Firstly, a small but significant number (94) appeared to be previously characterised interactions that had simply escaped annotation in APID (or by systematic interactome screens), based on the fact that the cross-linked proteins reside in the same CORUM-annotated complex (42) or even the same PDB entry. Secondly, 191 of the remaining novel PPIs were predicted with at least medium confidence by the STRING database (combined score of at least 0.4) (43), which integrates information across multiple lines of evidence including known interactions curated for homologous protein pairs, gene and protein co-expression, and literature text-mining. Thirdly, 676 of the remaining novel PPIs shared APID-annotated interaction partners and hence local interactomes (at one degree of separation).

Lastly, using the resources above, 197 PPIs of the novel PPIs remain ‘unexplained’. Of these, ~16% (32) involved the heat-shock protein Hspa1b (Uniprot: P0DMV9, (44)). Because the function of this protein involves promiscuous binding to many protein substrates, our cross-links might well be capturing biologically relevant interactions. In contrast, the CCT complex sub-interactome, which is known to be a more specific chaperone system, had many previously undetected PPIs but none that were categorised in the above scheme as unexplained.

### AlphaFold Multimer-v2 predicts structural interfaces for hundreds of cross-linked PPIs

One advantage of XL-MS over other systematic interactome mapping approaches is its ability to localise interaction interfaces rather than simply identify interactions. Because AF2 has recently been extended to predict the structures of protein complexes (AlphaFold Multimer) (45), we sought to use our data to inform and assess AF2-generated models of complexes. We chose the subset of 590 PPIs from our dataset that were captured by at least two URPs and subjected 530 of these (some interactions were too large to model by the software) to AlphaFold Multimer-v2 modelling (**Supplementary Table 5**). As a control, 400 randomly sampled protein pairs from the pool of proteins identified in our study were also modelled.

We assessed the overall quality of predicted structures of complexes by examining several structural measures. First, we examined the number of clashes within the models (identified by MolProbity (46)), which was reported to be unusually high (47) in the initial release of AlphaFold Multimer. However, most models had acceptably low levels of clashes (clashscore ≤ 100; *i.e*., less than 10% of atoms clashing) (**Supp Figure 7A**). We then used PISA (48) to identify structural interfaces in our models and defined ‘well-modelled’ interfaces as those that had an average pLDDT score of ≥ 70 for PISA-identified interface residues (**Supp Figure 7B**), a PISA-defined interface size of > 200 Å^2^ (**Supp Figure 7C**) and a clashscore ≤ 100. Using these criteria, AlphaFold Multimer-v2 generated at least one well-modelled structure (from the five generated per run) for a total of 343 of 530 (69%) cross-linked PPIs compared to 78 of the 400 (19%) random protein pairs (**Supplementary Table 5**).

Well-modelled AlphaFold Multimer-v2 models for PPIs defined by cross-links had significantly larger interaction surface areas than randomly sampled protein pairs (p < 2.2 × 10^-16^), corroborating the novel interactions defined by our data. Interestingly, the size of interaction interfaces also differed for PPIs that had previously been detected by different interactome mapping approaches. For example, PPIs described by binary interactome mapping approaches had, by comparing median values, 1.5-fold larger interface areas than those described only by indirect methods (AP-MS, BioID) (p = 0.003) (**Figure 4B**). Binary PPI interaction interfaces were also significantly larger than novel PPIs interfaces (>1.5-fold larger, p = 0.007). On the other hand, novel and known (but only) indirect PPIs did not significantly differ in interface size (p = 0.23). This observation perhaps flags differences in the nature of PPIs that can be detected by binary assays carried out in a heterologous system (*e.g*., Y2H) versus approaches (such as XL-MS or AP-MS) that retain native PTMs and additional complex partners. Hits in the former assays will be restricted to PPIs that are more robust and less dependent on biological context.

We were also able to use the experimental structures in the PDB to assess the AlphaFold Multimer-v2 dimer predictions. Of the 343 well-modelled complexes defined above, 151 comprised pairs of proteins that could be found in the same PDB entry. Furthermore, an additional 81 PPIs had PDB entries curated for homologous protein pairs (at least 40% sequence similarity). We superimposed the AlphaFold-derived structures onto their corresponding PDB structures for these 232 proteins and found that 154 pairs (66%) aligned well, with a median RMSD value of 1.6 Å (**Supplementary Table 5**).

In our dataset, AlphaFold Multimer-v2 performed most poorly in situations where the two proteins are part of the same complex but do not make direct contacts in the experimental structure. For example the Arp2/3 complex, which is involved in the formation of branched actin networks (49), comprises seven subunits and has had its structure determined by cryo-EM (PDB: 6UHC (50), **Figure 4C**). Based on the existence of cross-links, we predicted models for three subcomplexes: ARPC4-ARPC5 (Uniprot: P59998 and O15511), ARP2-ARPC2 (Uniprot IDs: P61160 and O15144), and ARP2-ARPC3 (Uniprot: P61160 and O15145). The prediction for ARPC4-ARPC5, which featured an interface of 830 Å^2^, closely matched the experimental structure (RMSD = 0.9 Å) and is supported by five URPs (**Figure 4C**). In contrast, although the other two subunit pairs do not make direct contact in the experimental structure, AlphaFold Multimer-v2 predicted spurious complexes that had interfaces of 830 and 430 Å^2^, respectively. All three URPs (two for ARP2-ARPC3, one for ARP2-ARPC2) did not fit the AlphaFold Multimer-v2 models for ARP2-ARPC2 (**Supp Figure 8A**) and ARP2-ARPC3 (**Supp Figure 8B**). In contrast, all three URPs were fulfilled in the experimental structure of the seven-subunit complex.

Similarly, we observed cross-links between pairs of histones such as histone H2A and histone H4. Although these subunits do make contact in the native nucleosome structure, they form more intimate dimers with H2B and H3 (PDB: 1AOI), respectively. However, because all four of these core histones have the same fold, AF incorrectly predicts a structure for the H2A-H4 complex that reflects the H3-H4 (or H2A-H2B) complex instead (**Supp Figure 8C**). It is noteworthy that misaligned histones alone account for 32 out of the 78 (41%) models that poorly match their experimental counterparts.

In summary, we find that although AlphaFold Multimer-v2 is largely correct in its model predictions, we would caution its use as a ‘discovery’ tool to identify PPIs. Such models should be corroborated with additional orthogonal experimental data such as XL-MS data.

### Cross-links provide new insight into AlphaFold Multimer-v2 complex models

We asked whether our URP distance restraints could experimentally corroborate the 343 well-modelled dimer interfaces calculated above. These dimers each feature between two and 39 URPs; however, many URPs connect residues that were not confidently modelled by AlphaFold Multimer-v2. We therefore used only URPs involving high-confidence residues (*i.e*., pLDDT ≥ 70 for both cross-linked residues) for model assessment. As a result, 97 models did not have any URPs that met this criterion, leaving 246 models that were described by 1–32 URPs (**Supplementary Table 5**). For comparison, we also simulated random URPs for each model. We found that the Euclidean distances measured for our experimental inter-protein URPs were considerably shorter than the randomly sampled URPs, and also shorter than the randomly sampled URPs in the 400 randomly sampled protein pairs (**Supp Figure 9**, p < 2.2 × 10^-16^ for both comparisons). Thus, our experimental cross-links are enriched at the interface of the predicted models, consistent with AlphaFold Multimer-v2 generating predictions that reflect the true structures of these complexes.

When assessing the overall degree of cross-link satisfaction in our well-modelled PPIs, we found that 167 out of 246 PPIs (68%) had a 100% URP satisfaction rate in at least one of the five AlphaFold Multimer-v2 models (**Figure 4D**), and 106 of those 167 had 100% URP satisfaction for all five models (**Figure 4E**). Interestingly, the distribution of cross-link satisfaction rates for dimer predictions was clearly bimodal, which we did not observe for monomeric predictions **(Supp Figure 6A)**. This suggests that AlphaFold Multimer-v2 was either getting the models largely right or largely wrong. It is possible that ‘wrong’ models arise from AlphaFold Multimer-v2 not having all the necessary information (e.g. additional interaction partners, PTMs). In summary, it further highlights the need for experimental data, such as XL-MS, to validate AlphaFold Multimer-v2 generated models.

Of the 246 PPIs with at least 1 high-confidence URP, 171 had either a pre-existing PDB structure or a PDB structure involving homologous protein pairs. As described above, superimposition of AlphaFold Multimer-v2 predictions and these experimental structures revealed that there were 129 PPI models that aligned well and 42 that did not align well. Strikingly, 83% (107 of 129) of the well-aligned models had 100% URP satisfaction rates, compared to just 60% (25 of 42) for the poorly aligned models. This difference between the two groups is increased further to 81% (well-aligned models) versus 46% (poorly aligned models) when histones were excluded (on the basis that histones are small proteins, meaning that cross-links are much more likely to be satisfied regardless of how the proteins were arranged relative to each other). This suggests that high confidence URPs could be used to discriminate between a correct well-modelled AlphaFold model and an incorrect well-modelled AlphaFold model.

Similar to the situation for monomeric AF2 models **(Figure 3D, E)**, some dimer models confirmed and extended existing experimental structures of human PPIs. For example, although no PDB structure exists for the human beta-actin ACTB (Uniprot: G5E9R0) in complex with the adenyl cyclase-associated protein 1 CAP1 (Uniprot: Q01518), there are two PDB entries (6FM2 and 6RSW (51, 52)) comprising the close homologues alpha-actin ACTA from rabbit (Uniprot: P68135) and CAP1 from mouse (Uniprot: P40124). Our AlphaFold Multimer-v2 model was both able to confirm conservation of the overall structure in the human ACTB-CAP1 complex and extend coverage of the interface itself via two corroborating inter-protein URPs (**Figure 5A**). This previously unresolved and cross-link validated interface indicates that the interaction between ACTB and the CAP1 protein is more intimate than previously appreciated, featuring 30 hydrogen bonds, 20 salt bridges and an interface area of 3,590 Å^2^ in our AF model versus a combined 19 hydrogen bonds, 17 salt bridges and an interface area 2,140 Å^2^ in the experimental X-ray crystal structures.

**Figure 5.**
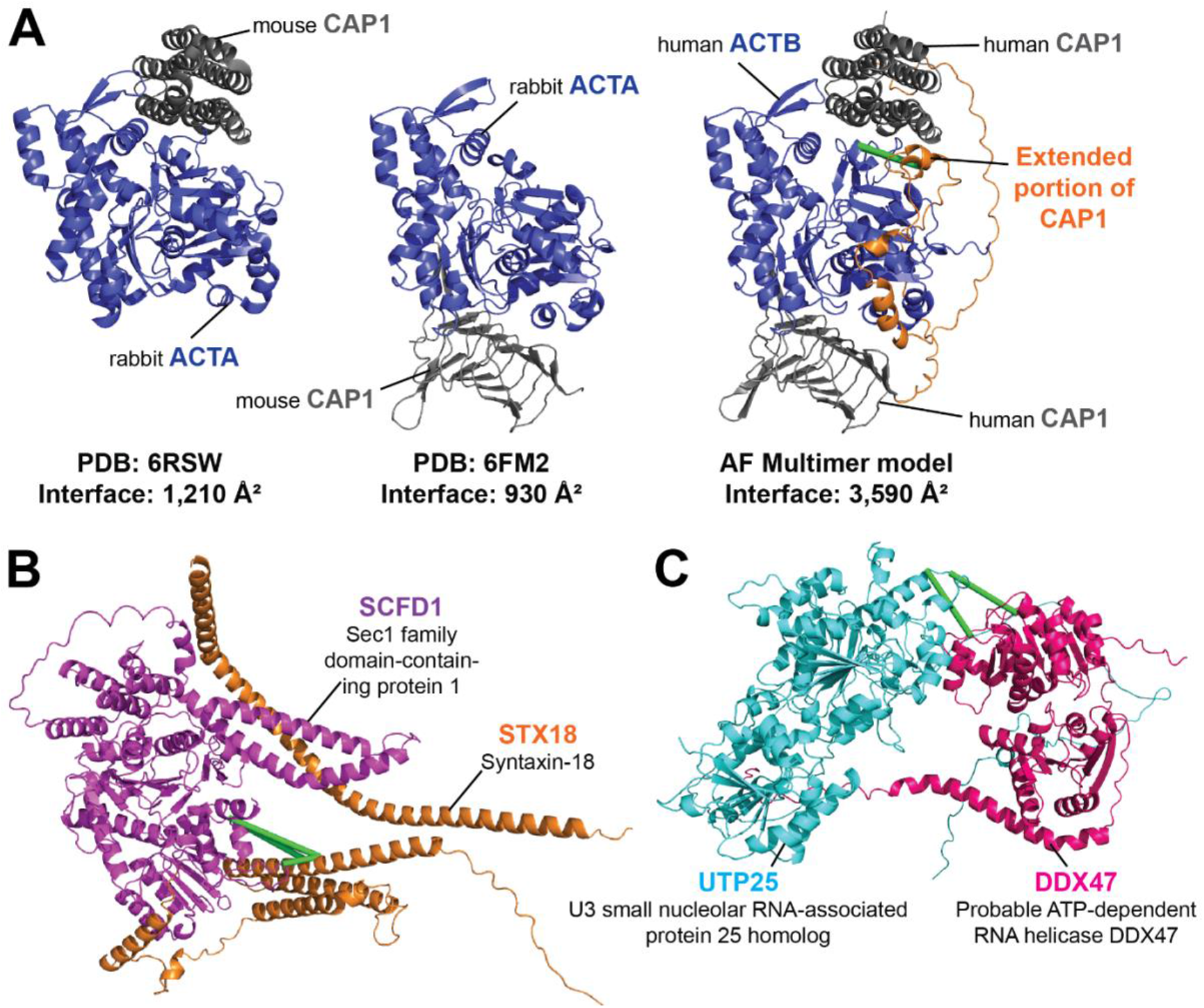
Cross-links enable assessment of computational predictions of structural interfaces mediating protein-protein interactions. **(A)** AF Mulitmer-v2 model of human beta actin ACTB in complex with the adenyl cyclase-associated protein 1 CAP1 (*middle*) versus the PDB structures 6FM2 (*left*) and 6RSW (*right*) of homologous proteins alpha actin ACTA from rabbit in complex with the mouse CAP1 protein. The AF multimer model largely recapitulates the regions resolved in the PDB structures and extends the interaction surface further (*orange*). **(B and C)** Examples of AF Multimer-v2 models of structurally undefined PPIs that have satisfied all URPs. Green lines denote satisfied URPs. **(B)** The model of the interaction between STX18 (Q9P2W9; *orange*) and SCFD1 (Q8WVM8; *magenta*). Four inter-protein URPs support this model. **(C)** The model of the interaction between UTP25 (Q68CQ4; *cyan*) and DDX47 (Q9H0S4; *pink*). Two inter-protein URPs support this model.

From our analyses, we also identified 75 PPIs with well-modelled AlphaFold Multimer-v2 interfaces that were structurally undefined (*i.e*., have no shared or homologous PDB entries available). Many of these PPIs had been previously characterised by binary and direct interaction mapping approaches. Of these 75 PPIs, 35 models had 100% URP satisfaction; these 35 models thus have a high likelihood of being correct. For example, the interaction of STX18 (Uniprot: Q9P2W9) and SCFD1 (Uniprot: Q8WVM8) was previously described by four studies (53–56). Our four inter-protein URPs were satisfied in all five models, and the top model featured an interface area of 3,688 Å^2^ with 21 hydrogen bonds and 15 salt bridges (**Figure 5B**). The formation of this PPI is implicated in the regulation of skeletal development (57) and, interestingly, there is evidence for an amyotrophic lateral sclerosis disease-causing missense mutation involving amino acid I70 in SCFD1 (I70T, ClinVar accession RCV001260556.1), a residue that is near cross-linked residues K63 and K61 at the interface. The combination of our data with AF thus provides precise and accurate structural context for biologically relevant PPIs.

Our data can also be used to both define new PPIs and experimentally corroborate their predicted interaction interface. For example, the top ranked model for the proposed interaction between UTP25 (Uniprot: Q68CQ4) and DDX47 (Uniprot: Q9H0S4) displays a 1,030 Å^2^ interface area and features two satisfied URPs (one of high confidence) (**Figure 5C**). The interface area and number of hydrogen bonds and salt bridges contributing to the interface (two for both categories) are significantly smaller than those seen in the previous two modelled PPIs described above. However, the two proteins were confidently predicted to interact in the STRING database (combined score of 0.98 out of a maximum of 1) and appear to have functional overlap. DDX47 is a DEAD-box RNA helicase known to associate with pre-RNAs (58) and UTP25 is implicated in pre-ribosomal rRNA processing (59, 60). Notably, both URPs defining this PPI were detected in the nuclear fraction, consistent with a shared role in ribosome biogenesis. It is possible that the presence of RNA cofactors or other protein complex partners could be required to stabilise this protein-protein interaction. This again highlights the ability of XL-MS to capture weaker interactions than some other interactome screening technologies and the value of the native context provided in our XL-MS approach.

### Cross-links define the binary interactions that underlie the assembly of larger protein complexes

Because our URPs can also be derived from higher-order assemblies, we mapped our data onto the CORUM database of manually-curated protein complexes (42). Our URPs could be mapped onto 366 unique CORUM-documented complexes (**Supplementary Table 6**). Of these 366 complexes, 165 displayed XLs for at least two distinct pairs of subunits. We considered the most densely cross-linked CORUM complexes, where ≥ 70% of proteins annotated in the complex were involved in at least one cross-linked protein-protein interaction. In total, there were 51 such complexes. Some, such as the CCT complex (**Supplementary Table 6**), are found in the PDB, with all cross-linked PPIs sharing PDB entries and 88% of the mappable URPs satisfied. However, almost half (21) of the densely cross-linked complexes were missing structural information for at least one PPI for which we found cross-linking evidence. These complexes could benefit from integrative structural modelling approaches, such as the cross-link guided modelling pipelines used in Assembline (61) and IMProv (62).

Additionally, we asked whether AlphaFold Multimer-v2 could be used to generate structures for higher order complexes and whether our cross-links could corroborate the models. The pentameric tRNA ligase complex comprising DDX1, FAM98B, RTRAF, RTCB and ASHWIN (Uniprot: Q92499, Q52LJ0, Q9Y224, Q9Y3I0 and Q9BVC5; CORUM #6301) plays an essential role in tRNA splicing (63). While there are structures (including homologous structures) of some individual subunits and domains (PDB: 7P3B (64); PDB: 7P3A (64); PDB: 4XW3 (65); PDB: 6O5F (66)), no structures of subcomplexes or the complete complex are available. Although AlphaFold Multimer-v2 failed to produce a model of the full complex, it successfully predicted a model for a core four-membered complex, based on existing biochemical data that showed that only DDX1, FAM98B, RTRAF and RTCB are essential for complex formation (64) (**Figure 6A**).

**Figure 6.**
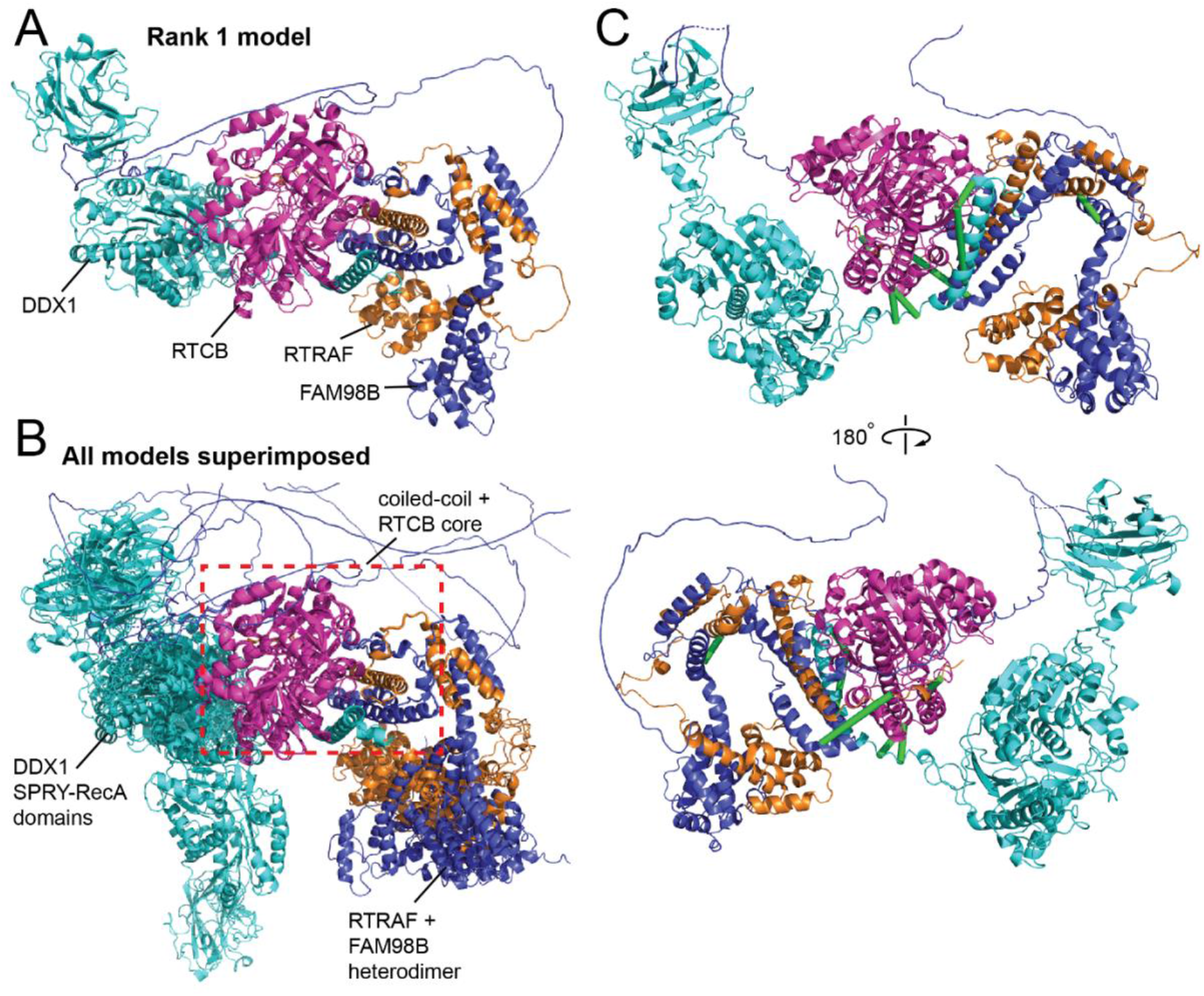
Cross-links support the AF Multimer model of the tRNA ligase core complex. DDX1 (Q92499) is in cyan, RTCB (Q9Y3I0) is in magenta, RTRAF (Q9Y224) is in orange and FAM98B (Q52LJ0) is in blue. **(A)** Rank 1 AF Multimer-v2 model of the tRNA ligase core complex. **(B)** All five AF Multimer-v2 models superimposed onto the Rank 1 model using the coiled-coiled core as the reference point. The coiled-coil core comprising of the C-terminal helices of DDX1, RTRAF and FAM98B together with RTCB is consistent across all five models (*red dashed box*). **(C)** Twelve URPs mapped onto the Rank 1 model of the tRNA ligase core complex. Eight URPs are fulfilled (*green lines*) The remaining four overlength URPs (*not shown*) were cross-linked to mobile regions of the DDX1 SPRY-RecA domains and the disordered C-terminal tail of RTRAF.

All five AF Multimer models featured a central helical bundle comprising the C-terminal helices of DDX1, FAM98B and RTRAF; this bundle packs against the RTCB ligase. In addition, the N-terminal domains of FAM98B and RTRAF formed an intimate heterodimer. Importantly, both observations are supported by available biochemical data (64). While the helical core of the complex was conserved in all models (superimposed RMSDs < 0.5 Å), the orientations of the SPRY-RecA domains of DDX1 and the FAM98B-RTRAF heterodimer showed considerable variability relative to the core (**Figure 6B**). Of the 12 URPs we measured, eight were fulfilled in all models (**Figure 6C**). Of the remaining four cross-links, two were to the disordered C-terminal tail of RTRAF and two were to the ‘mobile’ SPRY-RecA domains of DDX1. We had URPs between all members of the complex except to ASHWIN. In summary, our cross-links support the overall arrangement of the core of the complex that AF Multimer has predicted.

### The MIC60-MIC25-MIC19 hetero-tetramer has a 2:1:1 stoichiometry

Finally, we examined the seven-membered MICOS complex, for which we observed 12 URPs between five of the subunits. MICOS comprises MIC60, MIC13, MIC27, MIC25, MIC10, MIC26 and MIC19 (Uniprot: Q16891, Q5XKP0, Q6UXV4, Q9BRQ6, Q5TGZ0, Q9BUR5 and Q9NX63; CORUM #6255) and is essential for the proper formation and maintenance of crista junctions in the mitochondrial inner membrane (67). Despite its importance, the only existing structures are of distant fungal homologues of the coiled-coil domain of MIC60 (PDB: 7PUZ (64)) and of a complex between the C-terminal mitofilin domain of MIC60 and the CHCH domain of MIC19 (PDB: 7PV1 and 7PV0 (64)).

AlphaFold Multimer-v2 models of the full heptameric complex predicted that most of the sequences did not form ‘traditional’ globular folds (**Supp Figure 10**). Nevertheless, these models shared a core architecture that place MIC60, MIC25 and MIC19 in the same general spatial arrangement (**Supp Figure 10**), in agreement with published biochemical data (67).

The interaction between the mitofilin domain of MIC60 and the CHCH domain of MIC19 was also recapitulated in two out of the five calculated models. The other four subunits (MIC10, MIC13, MIC26 and MIC27) did not form substantial interactions with this core. MIC26 and MIC27 are apolipoproteins and is it possible that their presence in the complex is lipid-dependent.

Given that (a) MIC60 has been reported to form homo-oligomers (64), (b) a crystal structure corroborates the predicted MIC60-MIC19 interaction (64), and (c) the predicted structures of MIC19 and MIC25 were highly similar, we also used AlphaFold Multimer-v2 to explore the possibility that these proteins form a higher order assembly. Of the many permutations that we tested, the most striking – and the one predicted with highest confidence – was a 2:1:1 MIC60-MIC19-MIC25 complex (**Figure 7**). This complex displayed uniformly high pLDDT scores across all subunits (average pLDDT = 77) (which were higher than those observed for other assemblies tested; average pLDDT ≤ 70) and comprised a MIC60 homodimer bound symmetrically to MIC19 and MIC25 subunits via the MIC60 mitofilin domains and MIC19/MIC25 CHCH domains. The MIC60-MIC19 and MIC60-MIC25 interfaces closely resemble the MIC60-MIC19 crystal structure (PDB: 7PV0; RMSDs < 0.7 Å). Furthermore, all six inter-protein URPs and 12 (of 13) intra-protein URPs involving high-confidence residues are satisfied in this architecture (**Figure 7**), underscoring the orthogonal power provided by XL-MS data.

**Figure 7.**
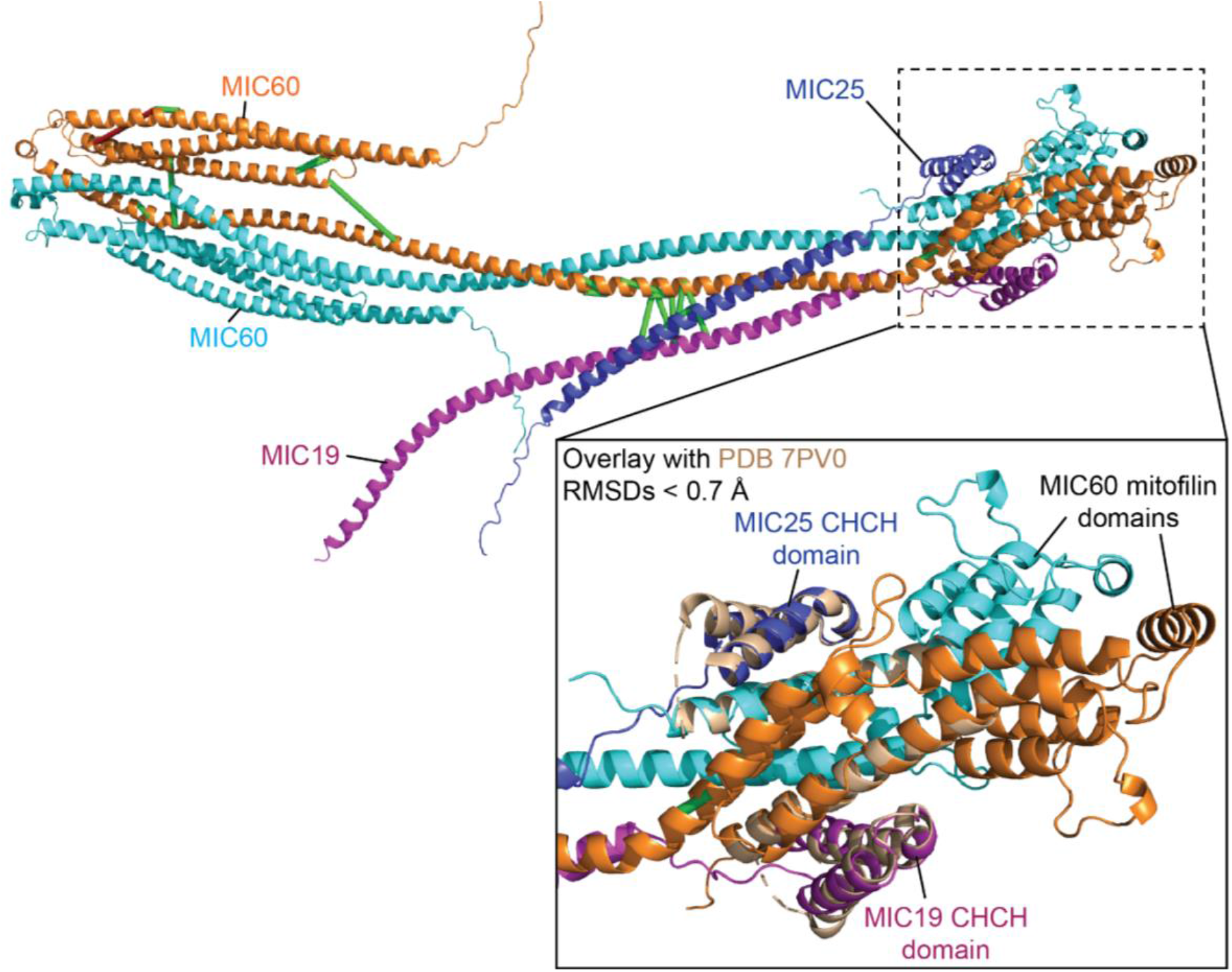
MIC60-MIC25-MIC19 is predicted to have a 2:1:1 stoichiometry. The representative rank 1 AF Multimer model of the 2:1:1 MIC60-MIC25-MIC19 subcomplex is shown. This complex displays uniformly high pLDDT score (average ~77) and comprises a MIC60 homodimer with its mitofilin domains bound symmetrically to the CHCH domains of MIC19 and MIC25. This MIC60-MIC19/MIC25 interaction overlays well (RMSDs < 0.7 Å) with the distant fungal MIC60-MIC19 structure (PDB: 7PV0; *biege*). MIC60 is in cyan and orange, MIC25 is in blue, MIC19 is in magenta.

In summary, we show that the combination of XL-MS data, AF Multimer-v2 and pre-existing biochemical data creates a powerful strategy for integrative modelling of higher order native protein complexes without the requirement for recombinant expression and purification.

## DISCUSSION

We have generated a deep cross-link resource reporting on more than 4,000 human proteins in near-native states with native sequences, abundances and contexts (*e.g*., the presence of native PTMs, relevant partner proteins as well as other macromolecules, cofactors and metabolites), which is a significant advantage over many other methods used to map protein structure and interactions. We have demonstrated that our cross-links are both of high quality and utility, and that they significantly extend and annotate the human structural proteome and interactome with very high throughput and biological relevance.

### Moving beyond the lysine cross-linked proteome

Our study highlights the value of moving beyond the lysine (K) to lysine (K) cross-linked proteome. Alternative cross-linker chemistries, such as bismaleimide (BMSO) (68) (homobifunctional, cysteines), can provide further, orthogonal information but have yet to be used routinely in proteome-wide studies. Photoreactive diazirine-based cross-linkers such as sulfosuccinimidyl 4,4’-azipentanoate (sulfo-SDA) (69), which allow cross-linking at any residue to be explored, represent a further extension. Although the greatest limitation for non-MS-cleavable cross-linkers of this type is the computing time required during peptide identification, the recent development of the MS-cleavable succinimidyl diazirine sulfoxide (SDASO) cross-linker (70) may aid in the feasibility and scale of such endeavours.

A recent community-wide, multi-lab study recommended that the search space for NHS-ester based cross-linking should be expanded to include S/T/Y residues (27). However, due to its prohibitive computational cost, searches including S/T/Y linkages are not routinely performed, especially in large-scale cross-linking studies. In our dataset and in line with previous conclusions (27, 28), a significant portion (~24%) of DSSO URPs involved S/T/Y residues. However, we note that only ~1% had S/T/Y to S/T/Y linkages, with most being K to S/T/Y linkages. This highlights that future studies should define NHS-ester reactivity as K to K/S/T/Y to balance computation time with localisation improvements and cross-link gains. This hybrid search strategy is currently configurable in several cross-link search engines such as pLink 2 (71) and MeroX (72). With a small compromise (reducing the size of the sequence database), we demonstrate that including S/T/Y linkages is possible on a large-scale basis.

Finally, although we have employed a diversified strategy to improve on the coverage of the cross-linked proteome using several cross-linkers combined with enrichment and fractionation strategies, many further possible enhancements exist. For example, the use of alternative proteases is relatively easy to implement but has seen limited use thus far (73, 74). Promising enzymes that will provide orthogonal information, include pepsin (cuts at Y/F/W), AspN (cuts at D) and the new ProAlanase enzyme (cuts at P/A) (75).

### A deep cross-linked proteome resource that complements AI-based structure predictions

We have demonstrated how our dataset can be leveraged to gain a deeper understanding of the biochemistry and biology of a higher eukaryote. Our structurally near-native cross-links corroborate experimentally determined protein structures – and with the advantage of being relatively fast, having modest sample requirements and providing data for proteins that are otherwise difficult to handle in isolation. We also identified regions of potential conformational variability and provide some evidence that PTMs might contribute to these variabilities. Furthermore, we show that XL-MS data can provide powerful corroboration of AI-based structure predictors like AlphaFold, which have democratised access to protein structure modelling for non-specialist groups and provide access to experimentally intractable proteins. With the recent release of more than 200 million predicted structures by AlphaFold (76) and 600 million in the ESM Metagenomic Atlas (77), it is now more important than ever to have orthogonal data to corroborate these models.

Perhaps most importantly, our data define 2,110 protein-protein interactions, a considerable extension in the coverage and diversity of the structural interactome. The URPs that define these PPIs can be used in combination with AI-based tools to generate and assess models of protein complexes, and such models can be subjected to experimental verification using biochemical and cellular approaches. Furthermore, regardless of AlphaFold modelling status, all 2,110 PPIs defined here contain cross-linking constraints that can help localise their interfaces at the sequence level without any reference to structure. Many interactions are known to occur exclusively in disordered regions, mediated by sequence motifs and domains (reviewed in (78)). Our cross-linking resource can therefore also be used to assist the annotation of domain-domain interactions and linear interaction motifs (as, for example, in Pfam (79)) - efforts that help to infer function for the thousands of understudied human proteins (12).

The wide accessibility of different cross-linkers, alongside critical efforts to standardise XL-MS search engines and analyses (80, 81), will enable high-volume and high-confidence cross-link constraints to be generated for proteins from diverse conditions and organisms. One could envisage a (near) future in which archived XL-MS data is harvested automatically by AlphaFold or similar systems and optionally displayed on predicted structures. Likewise, inter-protein cross-links could be displayed to alert the user that they could benefit from incorporating additional proteins into their prediction.

## METHODS

### Subcellular fractionation

The general procedure for subcellular fractionation were similar as described in (82, 83). For each biological replicate, 2 × 10^8^ freshly-harvested suspension HEK Expi293F™ (ThermoFisher Scientific) cells were used. Subcellular fractions were prepared by incubating the cell pellet with 5× the packed cell volume in swelling medium (15 mM KCl, 1.5 mM MgOAc, 10 mM HEPES-KOH, pH 7.5) for 10 min at 4°C. The cells were then centrifuged (300 × *g*, 2 min, 4°C) and the supernatant was discarded. The pellet was then homogenised by vortexing for 10 s, followed by repeatedly flushing the cells through a G-25 syringe. To this homogenate, 0.2 volumes of osmotic balancer buffer (375 mM KCl, 22.5 mM MgOAc, 1.25 M sucrose, 50 mM HEPES-KOH, pH 7.5) was added. The homogenate was then centrifuged (750 × *g*, 10 min, 4°C). This pellet consisted of the crude nuclei and was further treated as described below. The supernatant was transferred to a fresh tube and centrifuged at 3000 × *g*, 10 min, 4°C. This crude mitochondrial pellet was further treated as described below. The supernatant was again transferred to a fresh tube and centrifuged at 20,000 × *g*, 20 min, 4°C. The subsequent pellet, enriched with lysosomes, peroxisomes and Golgi membranes (henceforth described as the Golgi pellet for brevity), was further treated as described below. The supernatant should comprise of cytosolic and microsomal proteins.

The crude nuclear pellet was washed in resuspension buffer 1 (10 mM HEPES-NaOH, 0.25 M sucrose, 25 mM KCl, 5 mM MgCl2, pH 7.4) + 0.5% (v/v) IGEPAL CA-630 and recentrifuged (750 × *g*, 10 min, 4°C), twice. The final pellet was resuspended in resuspension buffer 1 (without IGEPAL CA-630). The crude mitochondrial pellet was washed in resuspension buffer 2 (10 mM HEPES-NaOH, 0.2 mM EDTA, 0.25 M sucrose, pH 7.4) and re-centrifuged (3000 × *g*, 10 min, 4°C), twice. The final pellet was resuspended in resuspension buffer 2. The Golgi pellet was washed in resuspension buffer 2 (10 mM HEPES-NaOH, 0.2 mM EDTA, 0.25 M sucrose, pH 7.4) and re-centrifuged (3000 × *g*, 10 min, 4°C) once. The final pellet was resuspended in resuspension buffer 2. Protein content of each subcellular fraction was quantified using a BCA protein assay kit (Pierce) and adjusted to 2 mg/mL prior to cross-linking.

### Protein cross-linking

For each cross-linking experiment, 2–6 mg of protein from each organellar fraction was used. The concentration of each cross-linker used for cross-linking experiments were pre-determined empirically, as in (84). Disuccinimidyl sulfoxide (DSSO; 100 mM stock solution in anhydrous DMSO) was added to a final concentration of 5 mM for all subcellular fractions except for the cytoplasmic fraction (1 mM instead) and allowed to react for 1 h. Dihydrazide sulfoxide (DHSO (22); 100 mM stock solution in Milli-Q water) in combination with 4-(4,6-dimethoxy-1,3,5-triazin-2-yl)-4-methylmorpholinium chloride (DMTMM; 200 mM stock solution in Milli-Q water) were added to final concentrations of 8 mM and 16 mM, respectively, and allowed to react for 1.5 h. Note: DMTMM is required for DHSO cross-linking, activating carboxyl groups in acidic residues to enable coupling with the hydrazide reactive groups in DHSO. DMTMM is itself also capable of directly catalysing carboxyl to primary amine coupling in amino acids residues to produce zero-length, but non-cleavable K-D/E cross-links. All reactions were incubated at 37°C.

Post-cross-linking, DSSO reactions were quenched with a final concentration of 100 mM NH4HCO3 at 37°C for 15 min, snap-frozen in liquid nitrogen and freeze-dried. As excess DHSO/DMTMM cannot be quenched, DHSO samples were chilled on ice, briefly sonicated to rupture organellar membranes, and 1 volume of ice-cold acetone was added. This mixture was then vortexed, and proteins were allowed to precipitate at −20°C for 2 h. Precipitated proteins were centrifuged at 20,000 × *g*, 15 min, 4°C. The supernatant was discarded, and the pellet was allowed to air-dry.

### Protein digestion and peptide fractionation

Cross-linked sample trypsinisation and peptide size exclusion chromatography were performed essentially as described previously (85). Briefly, dried, cross-linked samples were resuspended in 8 M urea to give a final concentration of 5 mg/mL of protein. Sonication was employed to help re-solubilise the protein pellets. Samples were then reduced (10 mM DTT, 37°C, 30 min) and alkylated (15 mM iodoacetamide, 20 min, room temperature in the dark). The samples were then diluted to 4 M urea with 50 mM Tris-HCl pH 8 and Trypsin/Lys-C mix (Promega) was added to an enzyme:substrate ratio of 1:250 (w/w) and incubated at 37°C, 4 h. Following this, the samples were further diluted to 0.8 M urea with 50 mM Tris-HCl pH 8, additional Trypsin (Promega) was added at an enzyme:substrate ratio of 1:200 (w/w), and the sample was further incubated at 37°C overnight (16 h minimum). After the overnight digestion, the samples were acidified with formic acid to 2% (v/v) and centrifuged at 16,000 × *g* for 10 min. The supernatant was then desalted using either 500-mg or 1-g Sep-Pak tC18 cartridges (Waters), and eluted in 60:40:0.1 acetonitrile:water:formic acid (v/v/v), snap-frozen in liquid nitrogen and freeze-dried.

For size exclusion chromatography fractionation (SEC), the dried desalted peptides were resuspended at 2 mg of peptide per 250 μL of SEC mobile phase (acetonitrile:water:trifluoroacetic acid, 30:70:0.1 (v/v/v)) and separated on a Superdex Peptide HR 10/30 column. A maximum of 2 mg of peptide was injected onto the column per SEC run. A flow rate of 0.5 mL/min was used and the separation was monitored by UV absorption at 215, 254 and 280 nm. Fractions were collected as 0.5-mL fractions. Based on the UV absorption traces, fractions of interest (retention volumes ~9–13 mL) were pooled, snap-frozen and freeze-dried.

Following SEC, peptides were further fractionated using high-pH reverse phase liquid chromatography (bRPLC). Dried peptides from the SEC step were resuspended at 2 mg per 4.5 mL of bRPLC buffer A (3% (v/v) acetonitrile, 5 mM ammonium formate, pH 8.5), filtered through 0.45 μm nylon filters and loaded onto a XBridge Peptide BEH C18 column (4.6 mm × 250 mm, 300 Å, 5 μm; Waters) using high pressure liquid chromatography (GBC LC1150) system. A maximum of 2 mg peptide was injected per run. Peptides were separated at a flow rate of 1 mL/min using a linear gradient of 3–40% bRPLC buffer B (80% (v/v) acetonitrile, 5 mM ammonium formate, pH 8.5) over 84 min, followed by a linear increase to 75% buffer B over 12 min. Peptides were monitored via UV (GBC LC1210) absorption at 215 and 280 nm. Ninety-six 1 mL fractions were collected across the whole bRPLC run. The collected fractions were then concatenated into 12 fractions by combining 8 fractions that are 12 fractions apart (*e.g*., the first pooled fraction comprised of original fractions 1, 13, 25, 37, 49, 61, 73 and 85) (86). The pooled fractions were then snap-frozen and freeze-dried.

### Mass spectrometry

Dried peptides were resuspended in 4% (v/v) acetonitrile, 0.1% (v/v) formic acid and loaded onto a 30 cm × 75 μm inner diameter column packed in-house with 1.9 μm C18AQ particles (Dr Maisch GmbH HPLC) using a Dionex UltiMate 3000 UHPLC (ThermoFisher Scientific). Peptides were separated using a linear gradient of 10–50% buffer B either over 81 min or 107 min at 300 nL/min at 55°C (buffer A consisted of 0.1% (v/v) formic acid, while buffer B was 80% (v/v) acetonitrile and 0.1% (v/v) formic acid).

Mass analyses were performed using either an Orbitrap Fusion tribrid or a Q-Exactive HF-X mass spectrometer (ThermoFisher Scientific). On the Orbitrap Fusion tribrid mass spectrometer, the 81-min gradient above was employed and the CID-EThcD-MS2-HCD-MS3 protocol (24) was used. Specifically, following each full-scan MS1 at 60,000 resolution at 200 *m/z* (350 – 1400 *m/z;* 50 ms injection time), precursor ions were selected for sequential CID-EThcD-MS2 acquisitions in a data-dependent manner (CID-MS2: *R* = 30,000, 18 NCE; EThcD-MS2: *R* = 30,000, calibrated charged dependent ETD parameters enabled, supplemental HCD at 30 NCE, 54 ms injection time; both: 1.6 *m/z* isolation window, 5 × 10^4^ intensity threshold, minimum charge state of +4, dynamic exclusion of 20 s). Subsequently, mass-difference-dependent HCD-MS3 acquisitions were triggered if a mass difference of (Δ = 31.9721 Da) was observed in the CID-MS2 spectrum (HCD, 30 NCE, 2 *m/z* isolation window; 5 × 10^3^ intensity threshold; charge state of +2–4; ion trap scan rate = rapid). Total duty cycle time = 1 s. Additionally, to maximise the detection of DMTMM-mediated cross-linked peptides, DHSO/DMTMM samples were also re-analysed on the Q-Exactive HF-X. For these analyses, the 107-min gradient above was employed and the following MS protocol was used: Following each full-scan MS1 at 60,000 resolution at 200 *m/z* (350 – 1400 *m/z*, AGC = 3 × 10^6^, 100 ms max injection time), up to 12 most abundant precursor ions were selected MS2 in a data-dependent manner (HCD, *R* = 15,000, AGC = 2 × 10^5^, stepped NCE = (25, 30, 35), 25 ms max injection time, 1.4 *m/z* isolation window, minimum charge state of +4; dynamic exclusion of 20 s).

### Identification of cross-linked peptides

DSSO and DHSO cross-linked peptides were identified using the XlinkX 2.0 (24) nodes as implemented in Proteome Discover v2.3 (ThermoFisher Scientific). The following key parameters were used: peptide mass between 300–10,000 Da, minimum peptide length of 5 residues, precursor mass tolerance ±10 ppm, product-ion mass tolerance of ±20 ppm for Orbitrap data and ±0.5 Da for ion trap data, allowable variable modification = oxidation (M), allowable static modification = carbamidomethyl (C), enzyme specificity of Trypsin with up to two missed cleavages (excluding the site of crosslinking), and FDR control of CSMs at 1%. The search database used was the UniProt human reference proteome (UP000005640; May 2020; 20,286 entries). For DSSO cross-links, the settings were as follows: allowable cross-linking sites were Lys, Ser, Tyr and Thr (or Lys-Lys only in pilot searches), cross-link mass-shift 158.0038 Da, cross-link fragment mass on cleavage = 54.0106 Da (alkene) and 85.9826 Da (thiol). For DHSO cross-links, the settings were as follows: allowable cross-linking sites were Asp and Glu, crosslink mass-shift 186.0575 Da, crosslink fragment mass on cleavage = 68.0375 Da (alkene) and 100.0095 Da (thiol). XlinkX score titrations were performed using the method described in (16) considering Lys-Lys linkages only, and analysing spectra from a smaller pilot experiment of DSSO cross-links (one biological replicate, four organellar fractions, 56 HPLC fractions – a total of 5,916 CSMs identified at default XlinkX settings). Three score combinations were assessed, where D = delta XlinkX score and S = XlinkX score: D4S40 (default), D10S60 (from (16)) and D20S80. Note that XlinkX FDR is approximated using the equation DD/(TT + DD) as XlinkX 2.3 does not report hybrid TD matches, where D = decoy database match and T = target database match. This can lead to inappropriate FDR control. The XlinkX score titration revealed that default score cut-offs (XlinkX score = 40, delta = 4; abbreviated as D4S40) produced inter-links with an inflated FDR even at the CSM level at which they are controlled, while D20S80 could produce CSMs of equivalent quality for known and novel protein-protein interactions, and control the false discovery rate to <1% at all levels of redundancy **(Supplementary Figure 1A and 1B)**. The D20S80 search setting was used for DSSO and DHSO spectra in the final dataset. Due to computational constraints, searches were done in batches of organelles and biological replicates. For DSSO, only Lys-Lys linkages were allowed in a first pass search against the whole human reference proteome (as described above), whilst in a second pass, all linkages (Lys, Ser, Tyr and Thr) were permitted using a reduced search database consisting of only the proteins identified in the first pass. The identifications resulting from these DSSO searches (D20S80, with Lys, Ser, Tyr and Thr specificity) were used for the final dataset.

DMTMM crosslinked peptides were identified using pLINK2 v2.3.9 (71). Key pLink 2 search parameters were as follows: Peptide mass between 600–10,000 Da and peptide length between 6–100 were considered, precursor mass tolerance ± 15 ppm, product-ion mass tolerance ± 20 ppm, variable modification = oxidation (M), fixed modification = carbamidomethyl (C), enzyme specificity of trypsin with up to two missed cleavages (excluding the site of cross-linking) per chain, and a 1% FDR. DMTMM cross-linker settings: cross-linking sites used in the final dataset were Asp, Glu and protein C-terminus to Lys and protein N-terminus, crosslink mass-shift –18.0106 Da. The search database used was the UniProt human reference proteome (UP000005640; May 2020; 20,286 entries). Decoy hits for decoy analysis were extracted from the unfiltered identification list table (the .csv output table with no suffix) by filtering for rows with the following attributes: Q.value <= 0.01, Peptide_Type = 3, Target_Decoy != 2 and isFilterIn = 1. We also considered the possibility of misidentified cross-linked peptides as co-eluted linear peptides (87) but did not find significant evidence of this occurring.

### Cross-link post-hoc filtering and consolidation

The R programming language (v4.1.3) was used to filter and consolidate DSSO, DHSO and DMTMM cross-link spectral matches (CSMs) across replicates and software outputs (XlinkX and pLink 2), and collapse redundant identifications to the levels of unique residue pairs (URPs) and protein-protein interactions (PPIs). For post-hoc filtering steps, decoy hits from XlinkX identifications of DSSO and DHSO data were removed. For pLink 2 identifications of DMTMM data, CSMs with a pLink score ≤ 0.34 were also filtered out to ensure appropriate FDR control at higher levels of redundancy (URP and PPI levels) **(Supplementary Figure 1C)**.

To consolidate filtered cross-links across software suites and replicates, the identity of each linked peptide’s sequence, the (peptide-based) site of cross-linker modification, and the linked amino acid type was parsed for each CSM from each software output. This enabled removing variability introduced across multiple pLink 2 and XlinkX search engine runs, where the assignment of master protein accessions (for peptide sequences redundantly mapped to the protein sequence database) can sometimes differ. Specifically, for each peptide sequence within the cross-link, the first protein accession (alphabetically) which mapped to the peptide was reassigned as the master protein for that sequence. This therefore standardised the protein(s) described by the CSM. The protein-level position of the cross-link modification in each peptide were then recalculated for use in the URP. If the two peptides could be mapped to the same protein sequence (an intra-protein cross-link), the first shared mapped protein accession was chosen to describe both proteins in the pair, with the protein-level positions recalculated accordingly for use in the URP. Homodimer URPs were defined when the sequences in an intra-protein cross-linked peptide overlapped.

### Biological annotations of proteins and protein-protein interactions

To annotate protein subcellular localisations, data was extracted from annotations of the human proteome hosted on the UniProt database (88). “Residents” of the isolated organellar fractions were those with relevant annotations for the “Subcellular Location” field or the cellular component Gene Ontology (89) term field were assessed for the presence of the following case-insensitive substrings; for the nuclear fraction (post-750 *g* pellet) – “nuclear”, “nucleus”, “histone”, “chromatin”, “nucleolus”, “nucleolar”, “nucleoplasm”; for the mitochondrial fraction (post-3,000 *g* pellet) – “mitochondria”, “mitochondrial”, “mitochondrion”, “mitoribosome”; for the Golgi (lysosome, perioxosome) fraction (post-20,000 *g* pellet) – ribosome”, “golgi”, “endoplasmic”, “reticulum”, “lysosome”, “peroxisome”; for the cytosol (microsome) fraction (supernatant post-20,000 *g*) – “cytosol”, “cytoplasm”, “microsome”.

To investigate protein abundances in the cross-linked proteome, the PaxDB protein abundance database (26) was used (downloaded August 2021). To annotate protein disorder, the MobiDB protein disorder database (37) (downloaded April 2022) was used. The aggregated majority consensus stringency was used (“prediction-disorder-th_50”) to determine whether the residues fell in disordered regions, which means at least 50% of the tested disorder predictor programs agreed on the annotation of the region as disordered.

To annotate the presence and density of PTMs within the sequences of cross-linked proteins, the PhosphoSitePlus database (36) (downloaded April 2022) of experimentally detected protein post-translational modifications (PTMs) was used.

The Agile Protein Interactomes DataServer (APID) database (2) (version 9606_noISI_Q3) of experimentally-derived protein-protein interactions (PPIs) was used to annotate the novelty and types of existing experimental evidence for cross-linked PPIs. The distinction of experimental interaction mapping techniques as either “binary” or “indirect” is used throughout this study and is based on the previous manual curations of experimental evidence codes defined by the original APID study and therefore annotated in their database (2). The resource was also used to determine the number of common interactors between two given proteins. The STRING-db resource (43) (v11.5) was used to determine the degree of predicted functional association between two proteins for random and cross-linked PPIs. Protein pairs without an association in the database were assigned a “combined score” of 0.

Only those with a “combined score” of at least 0.4 were considered functionally predicted. To simulate a random population of protein pairs, 300 protein pairs were randomly sampled from the cross-linked proteome. The CORUM database of protein complexes (42) (2018 release) was used to determine shared membership of a cross-linked PPI in a known protein complex. Only the first CORUM entry was considered in cases where the exact same group of proteins (including those that were not cross-linked) were annotated together in multiple complex entries.

### Protein structures

The Structure Integration with Function, Taxonomy and Sequence (SIFTS) database (3) (downloaded April 2022) was used to annotate experimental coverage of protein structures curated in the RCSB Protein Data Bank (4). This resource was also used to identify co-crystallised protein-protein interactions (two UniProt accession IDs present in the same PDB entry), and to determine the relevant PDB entries to download for cross-link mapping exercises. Structure files of interest (.pdb, asymmetric units) were downloaded in batch from the PDB database (4) website. For some large entries, the structures were only available as .cif files (due to the limit in number of atomic coordinates capable of being stored in the .pdb format). For these entries, the two linked chains of interest were extracted from the .cif files and then converted into .pdb format using the functions within the Bio3D R-package (v2.4-3) (90).

All AlphaFold predictions of monomeric protein structures used in this study were precomputed previously (9), and downloaded from the EBI AlphaFold Protein Structure Database (AlphaFold DB) (91) (v1 release). Only proteins which had models where their entire sequence length was considered in a single modelling run were analysed further (proteins < 2,700 amino acids).

Cross-linked PPI dimers (with at least 2 URPs) and random PPIs (sampled from the cross-linked proteome) were subjected to AlphaFold Multimer v2 modelling analyses. All modelled PPIs had a combined amino acid length of less than 2,000 residues. This was achieved by supplying canonical protein sequences for each complex member as the input for Colabfold (v1.2.0) (92), which uses MMSeqs2 (93) instead of JackHMMER (94) for accelerated MSA creation and implements AlphaFold-Multimer 2 (v2.2.0) (45). Colabfold was executed locally with the options --model-type AlphaFold2-multimer-v2 --recompile-all-models. The modelling with Colabfold was run on Nvidia Tesla Volta V100-SXM2-32GB GPUs on the Gadi supercomputer (National Computational Infrastructure, Australia).

For complexes with more than two members, the Google Collaboratory notebook version of ColabFold (v1.3.0) was used. The complexes were the pentameric tRNA ligase complex comprising DDX1, FAM98B, RTRAF, RTCB and ASHWIN (Uniprot: Q92499, Q52LJ0, Q9Y224, Q9Y3I0 and Q9BVC5) and the heptameric MICOS complex comprising MIC60, MIC13, MIC27, MIC25, MIC10, MIC26 and MIC19 (Uniprot: Q16891, Q5XKP0, Q6UXV4, Q9BRQ6, Q5TGZ0, Q9BUR5 and Q9NX63). For the tRNA ligase complex, two runs separate runs were performed comprising: (i) one copy each of all five full-length proteins; (ii) one copy each of full-length proteins of DDX1, FAM98B, RTRAF and RTCB. For the MICOS complex, four runs were performed comprising: (i) one copy each of all seven fulllength proteins; (ii) four copies of MIC60 residues (181–758); (iii) two copies of MIC60 residues (181–758), one copy of MIC19 residues (65–227) and one copy of MIC25 residues (110–235); (iv) two copies each of MIC60 residues (181–758), MIC19 residues (65–227) and MIC25 residues (110–235).

The command-line implementation of the Protein Interfaces, Surfaces and Assemblies (PISA) tool (v2.1.2) (48), as implemented in the CCP4 software suite (v4-7.1) (95), was used to calculate features of the structural interfaces predicted by the AlphaFold2 modelling. These included the number of detected inter-chain interfaces, the types and strength of chemical bonds, and the size and residues involved in the interface area.

To calculate the degree of atomic clashes in AlphaFold2 structural prediction models, the clashscore was calculated using MolProbity (46) as implemented in Phenix (v1.20.1-4487) (96), using the options o_flips=TRUE, coot=FALSE, probe_dots=FALSE. This was performed using the high-performance computing cluster hosted by the University of New South Wales (Katana). The pLDDT scores of individual residues in AlphaFold monomer models, and additionally the interface residues in dimer models identified by PISA, was accessed by extracting the b-factor variable in the .pdb file format using the Bio3d R-package (v2.4-3) (90). All protein structures were visualised using the PyMOL software (v2.5.2).

### Cross-link structure mapping

The Xwalk command line executable program (v0.6) (29) was used (with the -dist, -bb, -f and -euc flag options) to determine the Euclidean distances (alpha-carbon to alpha-carbon) between cross-linked residues on experimental or predicted protein structures previously downloaded (or converted into) .pdb files. For experimental structures, random URPs were generated for each PDB entry by sampling fifteen random but theoretically valid residue pairs (5 per cross-linker; KSTY-KSTY, K-DE, DE-DE) from the protein sequences present in the given PDB. For predicted monomeric structures, these were sampled from the modelled protein sequence. For predicted dimer structures, three random URPs (one for each cross-linker reactivity pair) were sampled where one residue was sampled from the first protein sequence and the other from the second. Xwalk input files were then generated for each structure containing both the experimental and random URPs. For experimental PDB files, this involved accounting for any PDB entry-specific differences in the residue index from that reported in the canonical protein sequences used for cross-link identification. This was achieved by applying a numeric adjustment to the cross-linked residue number by reference to the annotations in the SIFTS resource (3). In the case of chain ambiguity within multimer PDB structures, URPs were redundantly described so that each theoretically possible combination of chain pairs could be considered. For analyses where the span of distances across multiple PDB entries was calculated, only structures with unique chain mappings (no multimeric chains of the same protein) were used.

Cross-links were visualised on structures in Pymol (version 2.5.2), using either the automated .pml scripts generated by the Xwalk command line tool (version 0.6, using the flags -dist, -bb, -f, -euc, and -pymol) (29) or at a smaller scale, using the PyXlinkViewer PyMOL plugin (97).

### Homology search and structural alignments to PDB structures containing homologous proteins

To identify experimental structures existing for protein sequences homologous to cross-linked proteins, protein sequences were subjected to BLAST analysis (39) to search for homologous sequences. The protein sequences were downloaded in FASTA format by batch retrieval of UniProtKB IDs from the UniProt website. Either the webserver, or a command-line implementation of the NCBI BLAST+ blastp program constructed using the functions within the biopython Python-package (v1.79), was used to query these sequences against a locally downloaded ‘pdbaa’ database (retrieved from the NCBI FTP server). Protein homologues were defined to be any matches from this BLAST search pertaining to an e-value < 1 × 10^-50^. A set of all PDB structure identifiers containing a homologue was obtained for each protein. For protein pairs, the intersection between the individual sets of homologue-containing structures were taken for further analysis.

Structural alignment was performed for each protein pair that had homologue-containing PDB structures. For each pair, the structure with the greatest homology, as determined by the smallest e-values from the BLAST search, was obtained in .cif file format using functions from the biopython Python-package (v1.79) to access the wwPDB API. The relevant chains in the PDB entry, as specified in the BLAST homology search, were chosen and aligned to the AF Multimer structure using functions from the pymol Python-package (v2.5.4). Structural alignments were stored in .pdb format and RMSD values from the alignment were saved.

### Statistical Analyses

Data manipulation and statistical analyses were performed within R (v4.1.3) using RStudio, using the inbuilt statistical functions such as the Wilcoxon signed rank test with continuity correction “wilcox.test”.

## ADDITIONAL INFORMATION

### Data Availability

All mass spectrometry data and XLMS search results have been deposited to the ProteomeXchange Consortium via the PRIDE partner repository (98) with the dataset identifier PXD035844. The dimeric models are available in ModelArchive (modelarchive.org) with the accession codes ma-low-csi.

### Code Availability

Key data processing scripts (including for the filtering and consolidation of cross-linked spectral matches across software suites) have been uploaded to FigShare.

## Acknowledgements

We acknowledge the Sydney Mass Spectrometry Core Research Facility at the University of Sydney for providing access to mass spectrometers and thank the technical staff for the maintenance of the instruments. This project was undertaken with the assistance of resources and services from the National Computational Infrastructure (NCI), which is supported by the Australian Government. T.K. B. acknowledges the support of an Australian Government Research Training Program scholarship. M.R.W. acknowledges support from the Australian Research Council, Australia, and J.P.M acknowledges support from the National Health and Medical Research Council, Australia. M.R.W. and X.V.C. acknowledge support from the New South Wales State Government RAAP scheme and the UNSW RIS scheme. J.P.M. acknowledges support from the Australian National Health and Medical Research Council.

## Author contributions

Conceptualization, T.K.B., J.P.M., J.K.K.L.; Methodology, T.K.B., J.P.M., J.K.K.L.; Software, T.K.B., X.V.C., C.L.; Investigation, T.K.B., X.V.C., J.K.K.L.; Formal analysis, T.K.B., C.L., J.P.M., J.K.K.L.; Resources, A.N., R.J.P., M.R.W., J.P.M.; Writing (original draft) – T.K.B., J.P.M., J.K.K.L.; Writing (review and editing) – T.K.B., X.V.C., R.J.P., M.R.W., J.P.M., J.K.K.L.; Supervision, M.R.W., J.P.M., J.K.K.L.

## Materials & Correspondence

All correspondence should be directly to Dr. Jason Low.

