## Supplementary Figures for "Cross-linking mass spectrometry discovers, evaluates, and validates the experimental and predicted structural proteome"

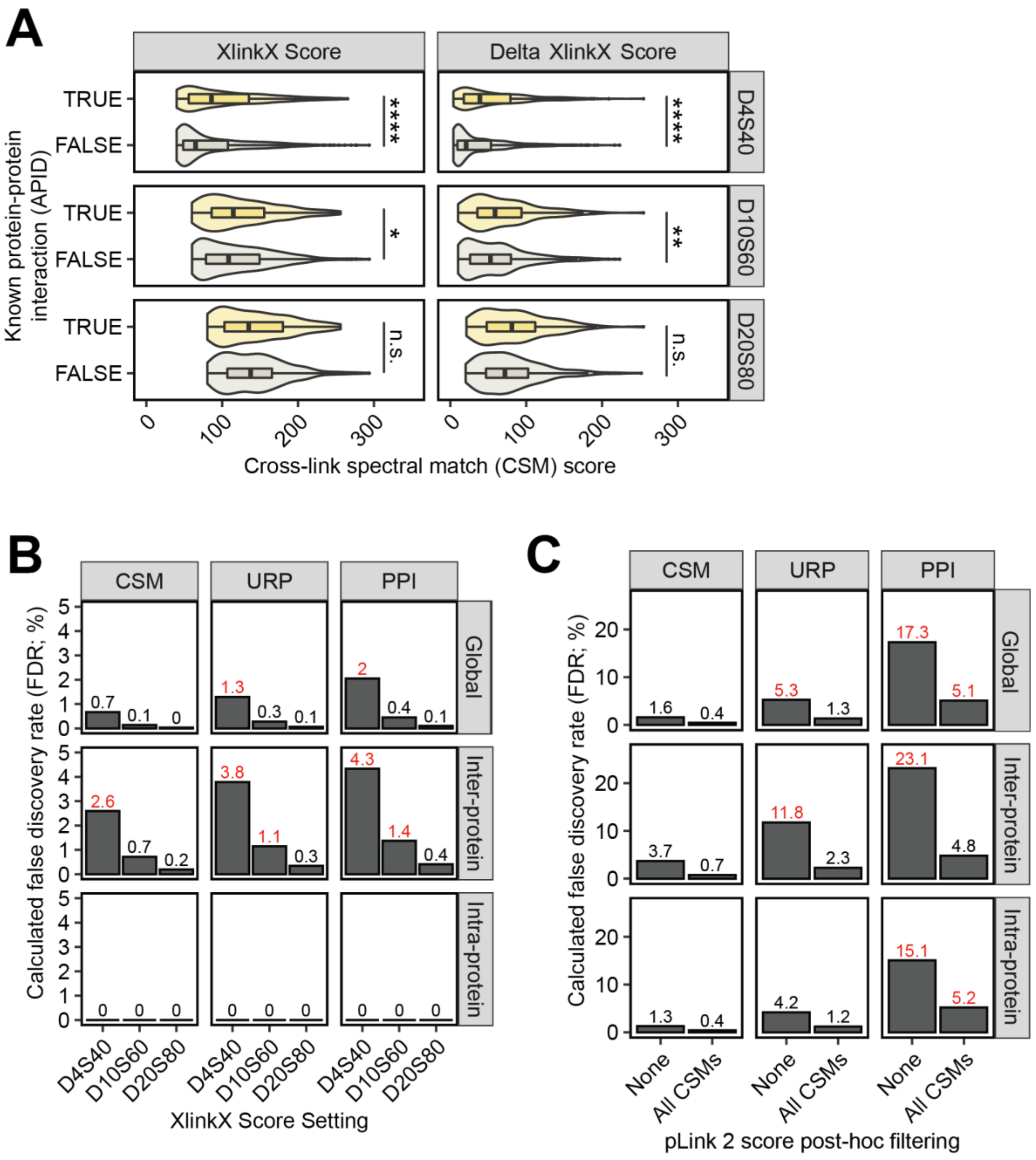


**Supp Figure 1: False discovery rate control measures to reduce incorrect identifications and propagation of error. (A)** and **(B)** show the results of XlinkX search settings titrated on a small pilot dataset, performed by varying XlinkX score cut-off settings in the XlinkX Search node. These search settings enable filtering of matched spectra (and importantly, individual peptide identifications within the cross-link) before FDR control by the XlinkX 'validator' node, which was set to 1% at the cross-link spectral match (CSM) level. The scores used were D4S40 (default settings, minimum delta XlinkX score of 4, minimum XlinkX score 40), D10S60 (as used in (1), minimum delta XlinkX score of 10, minimum XlinkX score 60) and D20S80 (as used for our final dataset, minimum delta XlinkX score of 20, minimum XlinkX score 80). **(A)** The distribution of inter-protein CSM XlinkX scores stratified by protein-interaction novelty as annotated by the APID PPI meta-database. This revealed that D20S80 produced CSMs of equivalent quality between known and novel protein-protein interactions (PPIs). Asterisks show results of Wilcoxon rank sum tests with continuity corrections, where

**** is $p < 0.0001$, ** is $p < 0.01$, * is $p \leq 0.05$ and ‘n.s.’ is $p > 0.05$. **(B)**, **(C)** Calculated false discovery rate (FDR) at different levels of biological interest (URP-level for intra-protein, and PPI-level for inter-protein links). Decoy cross-linked peptide sequences were compared to determine whether they originated from the same reverse protein sequence (intra) or two different sequences (inter) and used to calculate global, inter-link and intra-link false discovery rates at the CSM, URP and protein-pair level. **(B)** XlinkX score titration from the pilot datasets show that the D20S80 search setting was effective in controlling the FDR to < 1% at all levels of redundancy. **(C)** Post-hoc filtered pLink 2 DMTMM identifications from the full dataset. Note: although pLink 2 is reportedly quite effective at controlling FDR, we performed post-hoc score filtering to further control for the propagation of error at the URP and PPI-levels. CSMs were left as reported by pLink 2 (and after in-built FDR control) “None”, or filtered for having pLink scores ≤ 0.34 in inter-protein CSMs only or across all CSMs. The filtering strategy for pLink 2 CSMs chosen for our final dataset was to post-hoc filter only the inter-protein CSMs for scores ≤ 0.34, as this filtered intra-protein URPs and inter-protein protein-pairs to < 5% FDR.

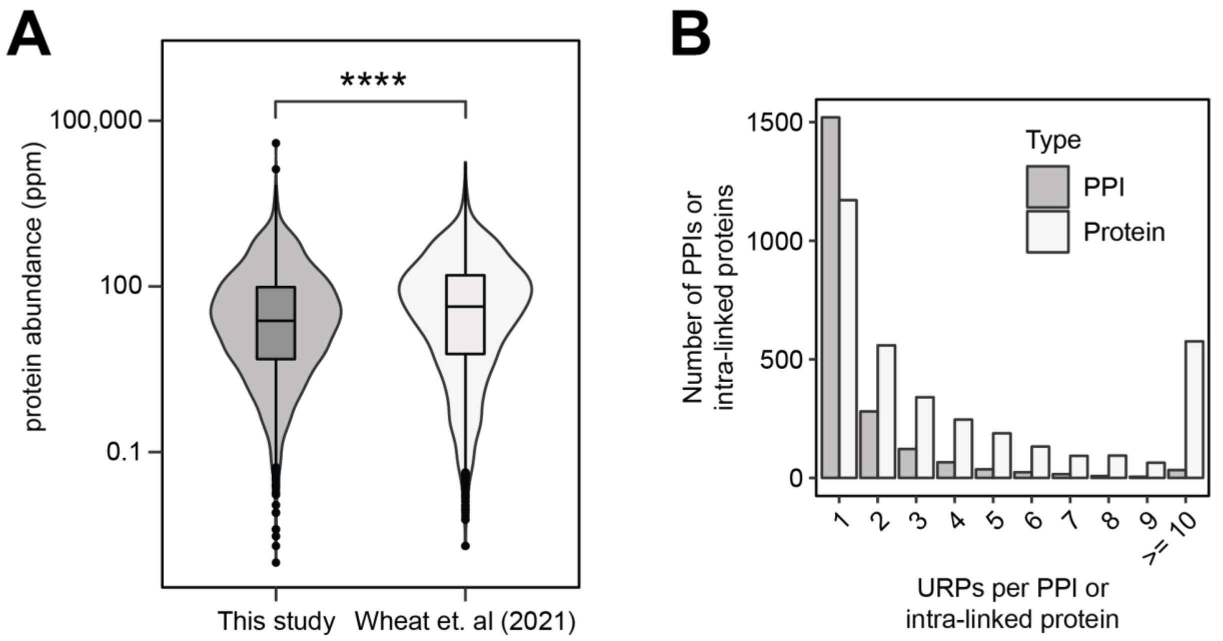


**Supp Figure 2. High density and deep coverage of the cross-linked proteome. (A)** PaxDB-annotated abundances for cross-linked proteins in this study and those identified in (2). **** = p < 2.2e-16 in a Wilcoxon rank sum test with continuity correction. **(B)** Histogram of URPs summarised per protein-protein interaction (inter-protein or homo-oligomeric URPs; *grey bars*), or per unique intra-linked protein (intra-protein URPs; *white bars*).

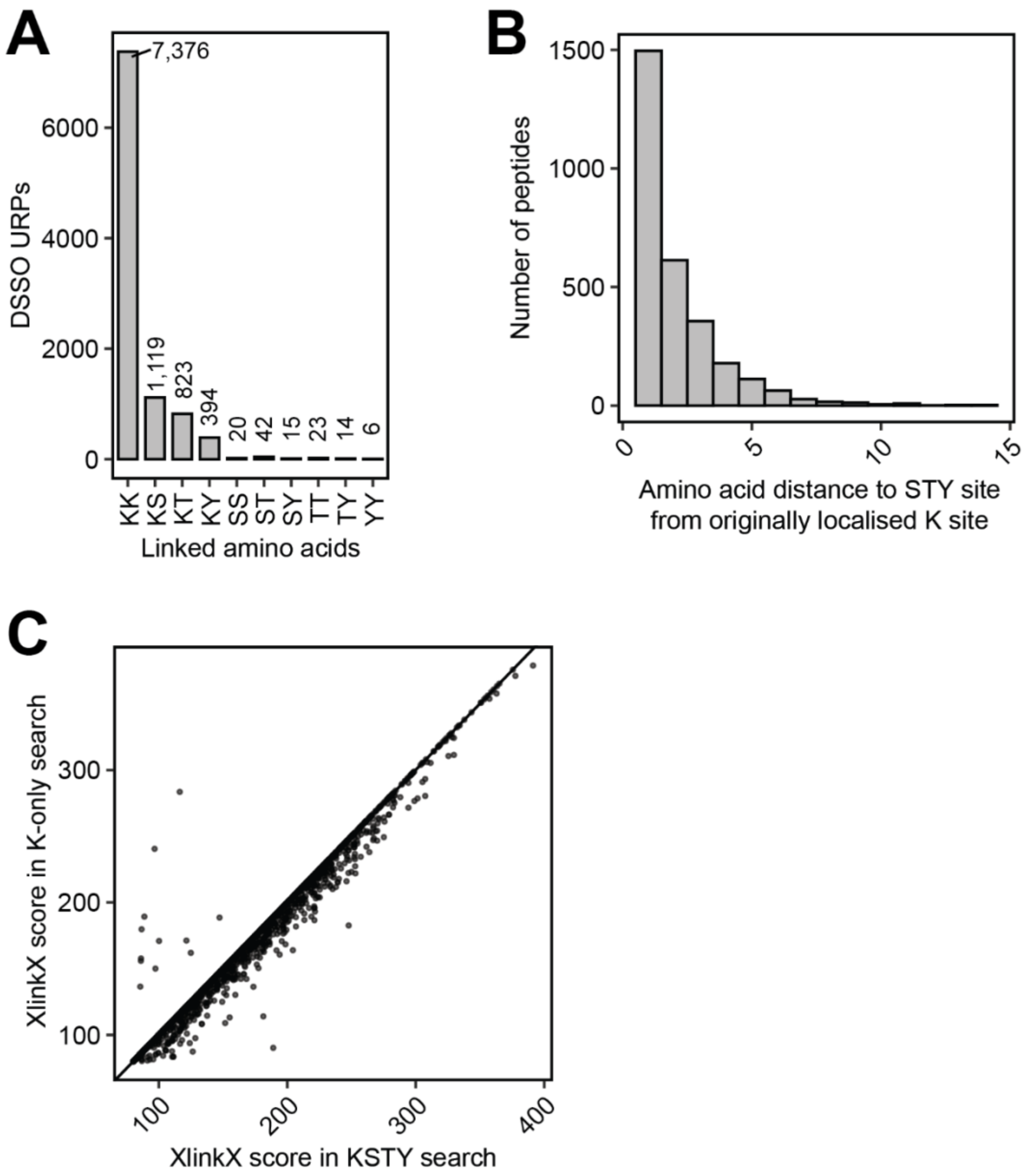


**Supp Figure 3. Consideration of DSSO S/T/Y reactivity improves quality of cross-links.** **(A)** The breakdown of amino-acid pairs in the final DSSO dataset. **(B)** Distance to nearest lysine-localised residue from K-K searches for peptides where the site of cross-link modification was reassigned as a S/T/Y residue. **(C)** The comparison of cross-link spectral quality scores for matched spectra containing peptides with differently localised cross-linker modified residues between searches are shown for DSSO cross-links identified with XlinkX**.**

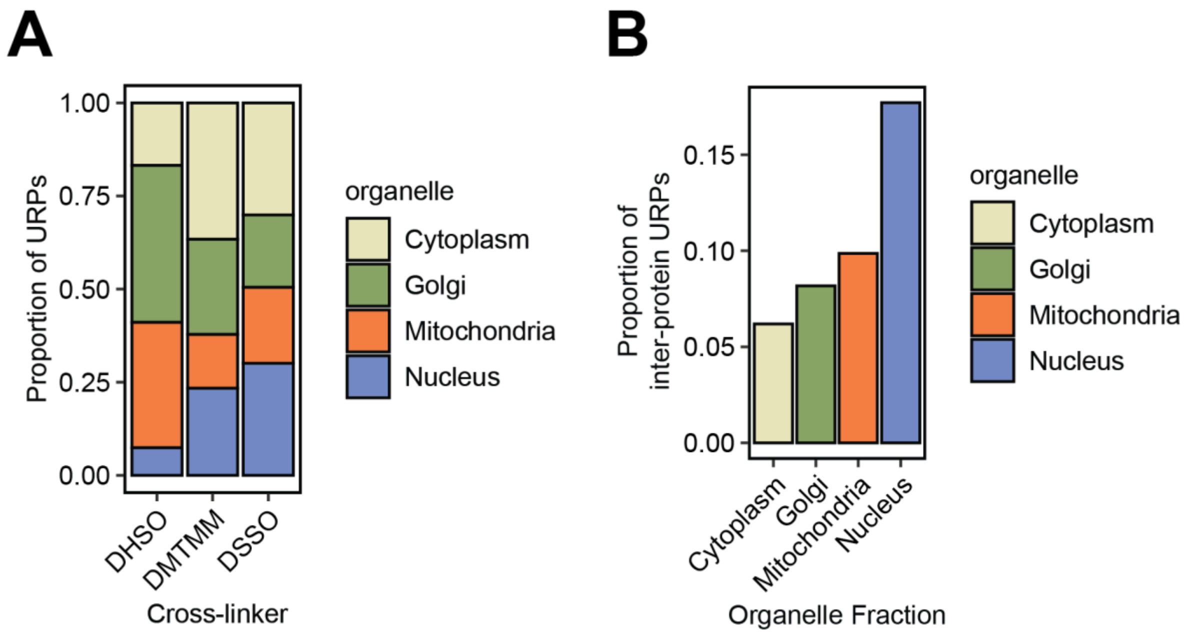


**Supp Figure 4. Effectiveness of each cross-linker in profiling different subcellular niches. (A)** The proportions of URPs identified from each organellar fraction differ between cross-linkers types. **(B)** The organelle fractions themselves differ in their proportion of identified inter-protein URPs. The definition of inter-protein in this graph includes unambiguous homo-dimer URPs.

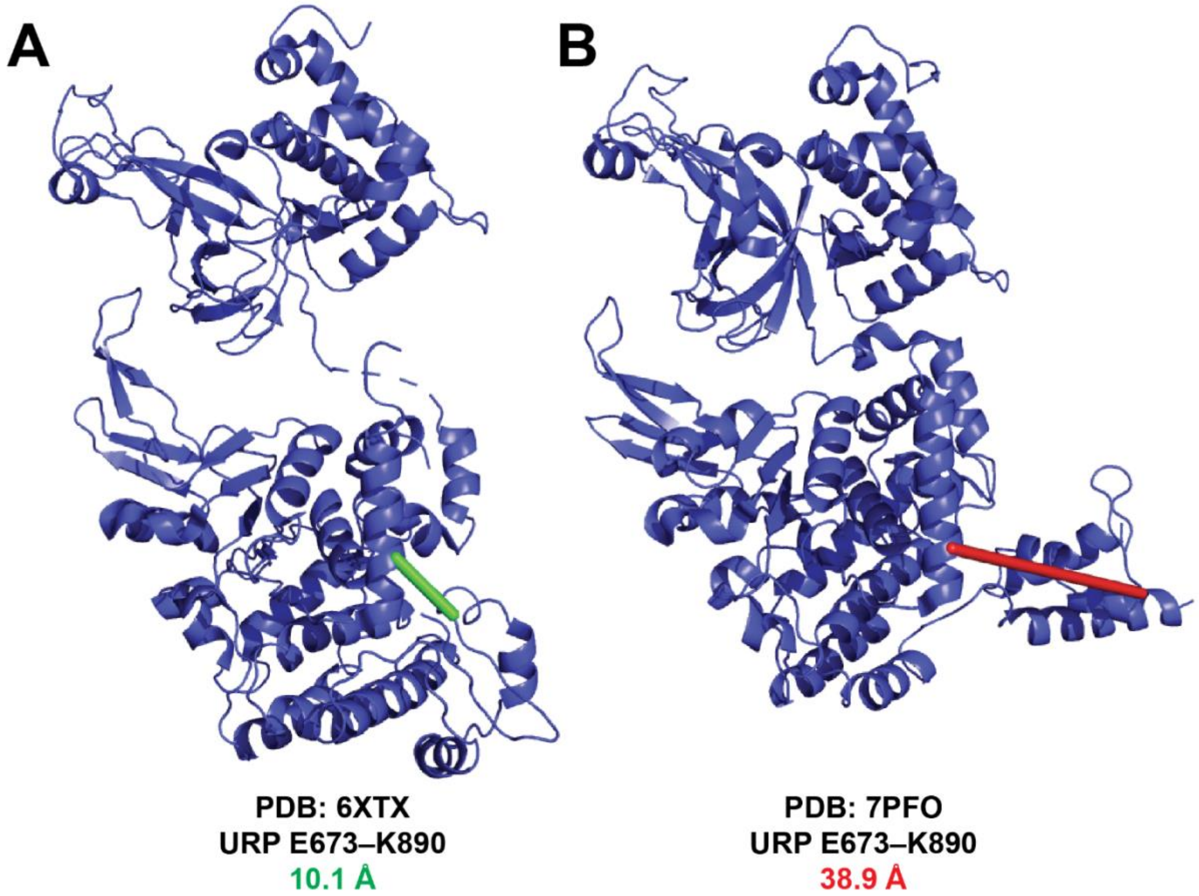


**Supp Figure 5. The URP E673-K890 in the DNA replication licensing factor MCM2 reports different cross-linker distances in different contexts.** MCM2 (P49736) is a member of the CDC45-MCM-GINS (CMG) helicase. **(A)** When the CMG helicase is not engaged in the replisome, the URP E673-K890 reports a distance of ~10 Å. **(B)** When the CMG helicase is engaged in the replisome, the URP E673-K890 reports a distance of ~39 Å.

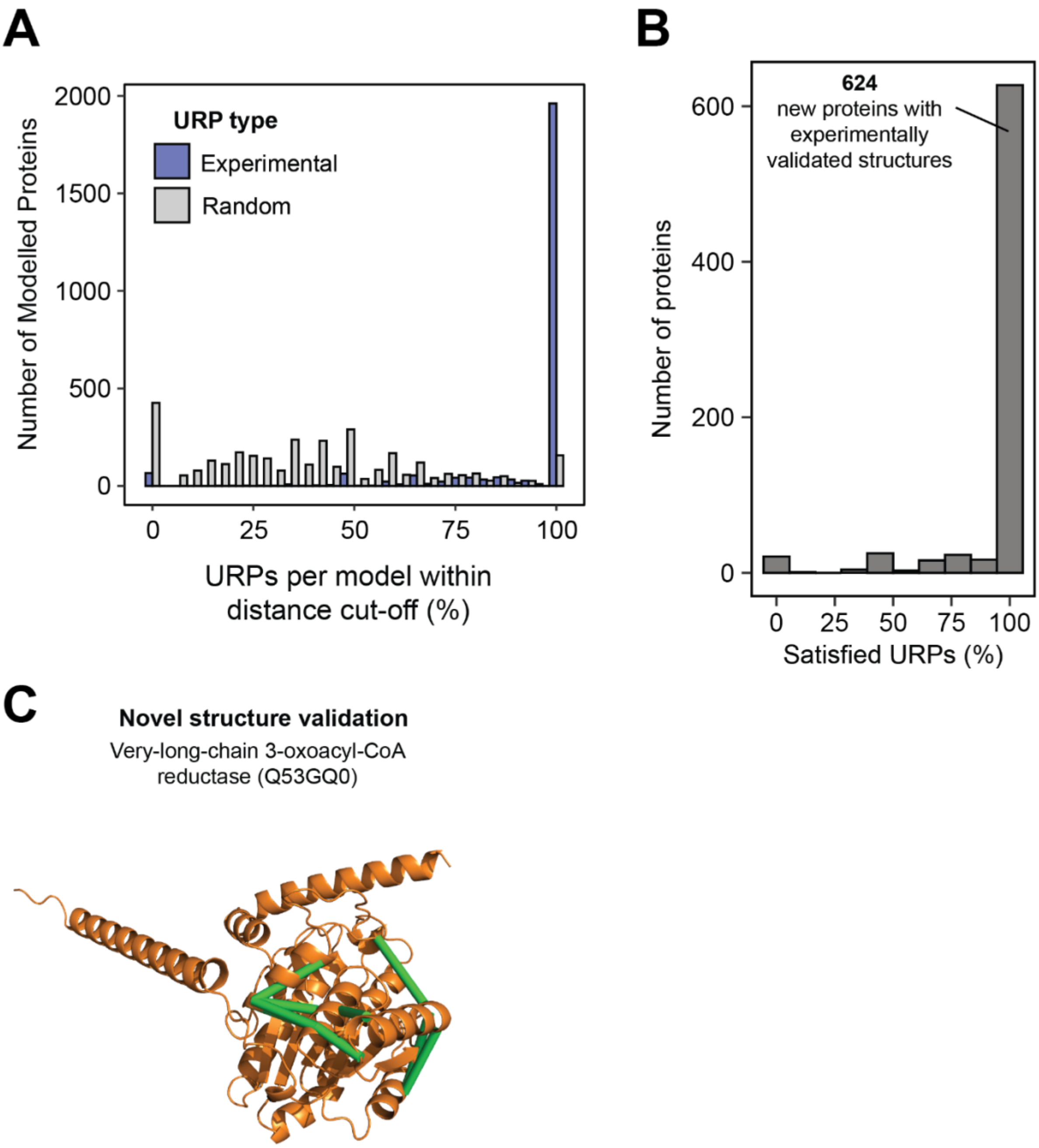


**Supp Figure 6. Cross-links validate AlphaFold2 monomer models of proteins absent from the PDB. (A)** The satisfaction rate for experimental and random URPs per AlphaFold2 model. For each model, five random URPs were generated for each cross-linker with appropriate sidechain reactivities. **(B)** 624 out of the 737 proteins without corresponding PDB entries satisfied all their high-confidence URPs in their corresponding AF2 models. **(C)** AlphaFold2 model of the very-long-chain 3-oxoacyl-CoA reductase enzyme (Q53GQ0) that does not have any PDB entries for itself or of homologous proteins. Nine URPs support the AF2 model.

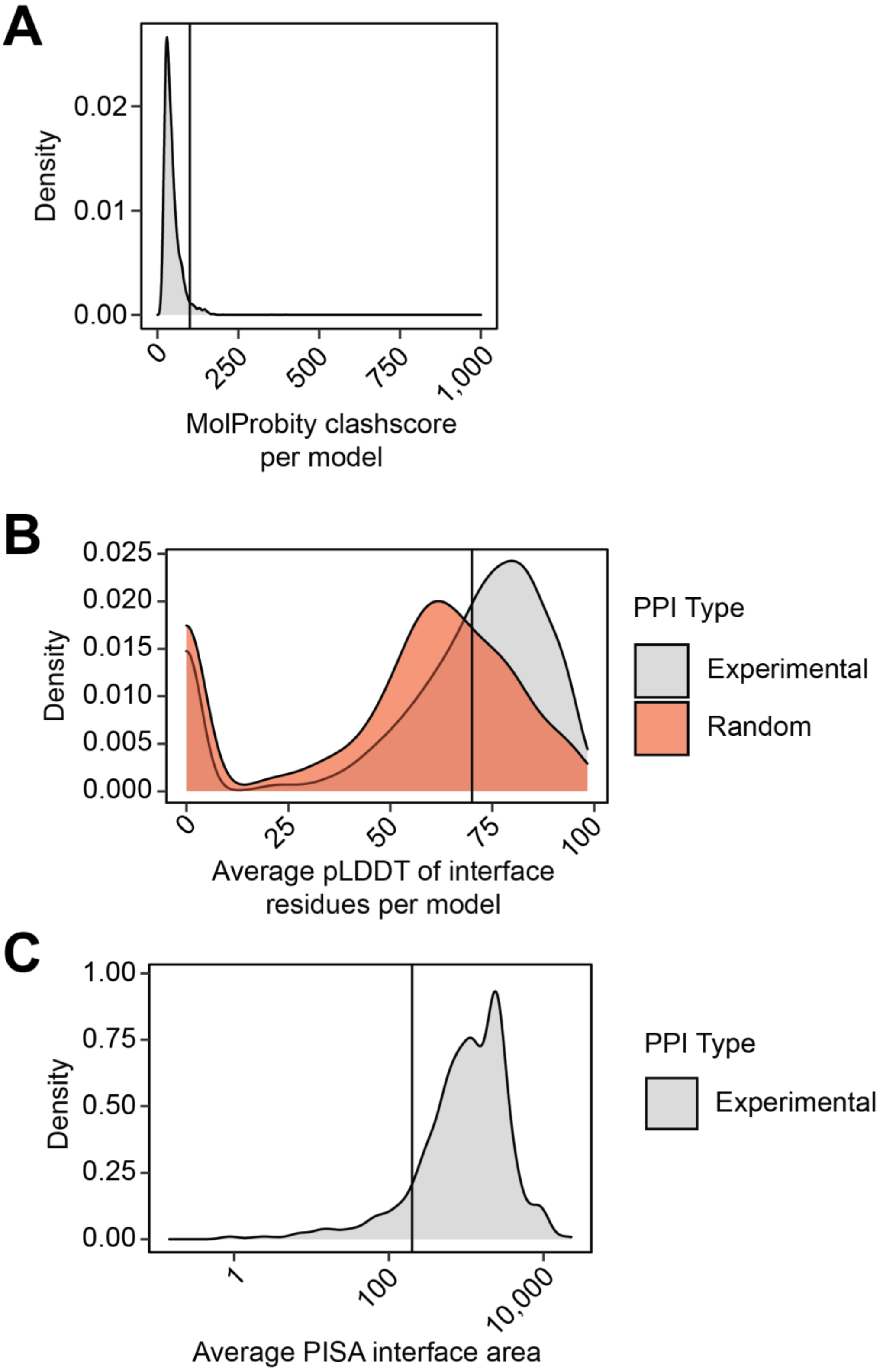


**Supp Figure 7. AlphaFold Multimer-v2 modelling quality statistics.** The distributions of **(A)** MolProbity clashscores (clashing atoms per 1000 atoms); and **(B)** the average interface residue pLDDT scores for all models generated in this analysis; and **(C)** the average interface area as defined by PISA. The line shows the cut-off used to filter for "well-modelled" interfaces.

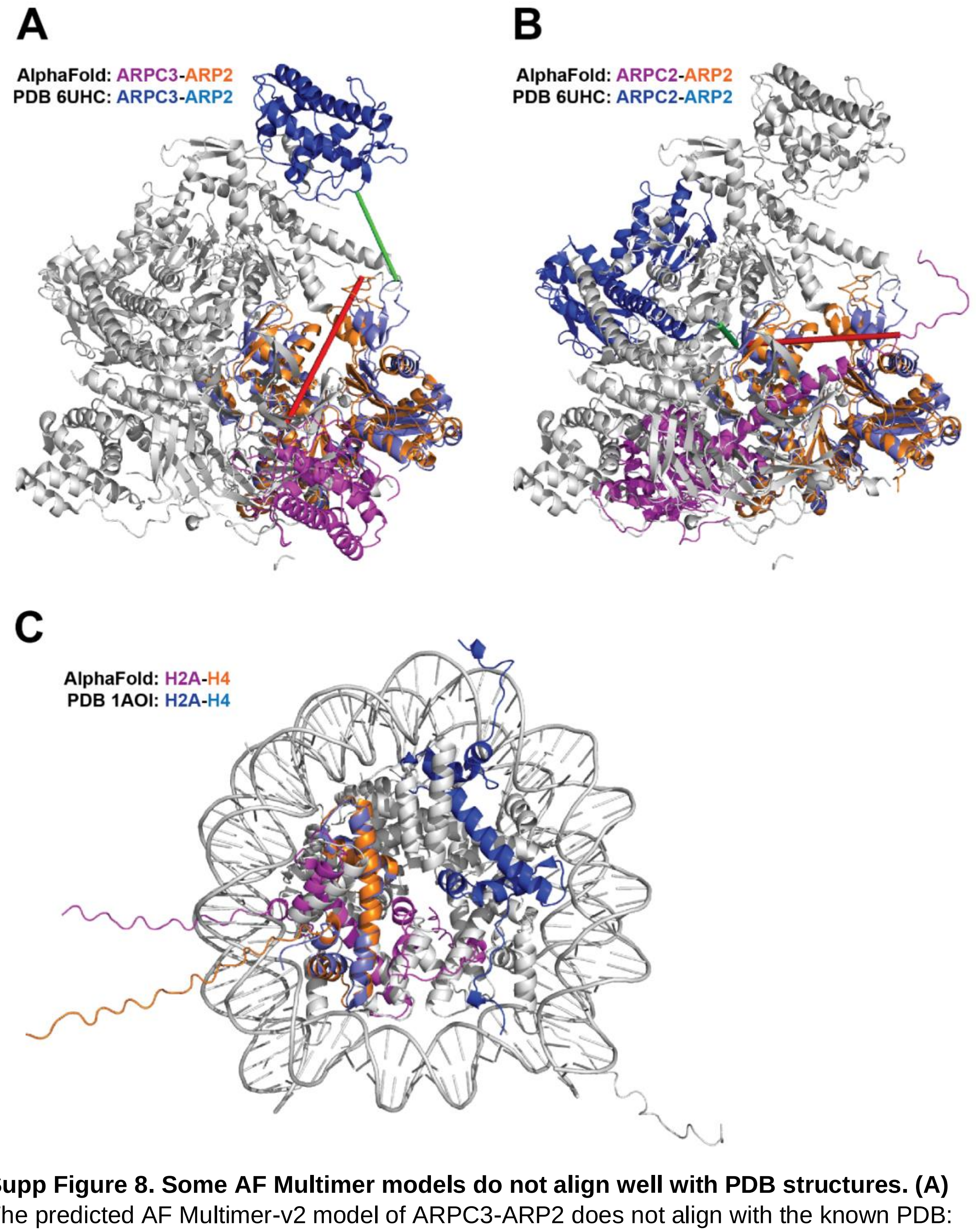


**Supp Figure 8. Some AF Multimer models do not align well with PDB structures. (A)** The predicted AF Multimer-v2 model of ARPC3-ARP2 does not align with the known PDB: 6UHC structure of the full 7-subunit Arp2/3 complex. Two URPs are available and both do not fit the AF model but are satisfied in the PDB structure. **(B)** The predicted AF Multimer-v2 model of ARPC2-ARP2 does not align with the known PDB: 6UHC structure of the full 7-subunit Arp2/3 complex. One URP is available and does not fit the AF model but is satisfied in the PDB structure. **(C)** The predicted AF Multimer-v2 model of histones H2A-H4 does not align with the known PDB: 1AOI structure of the nucleosome.

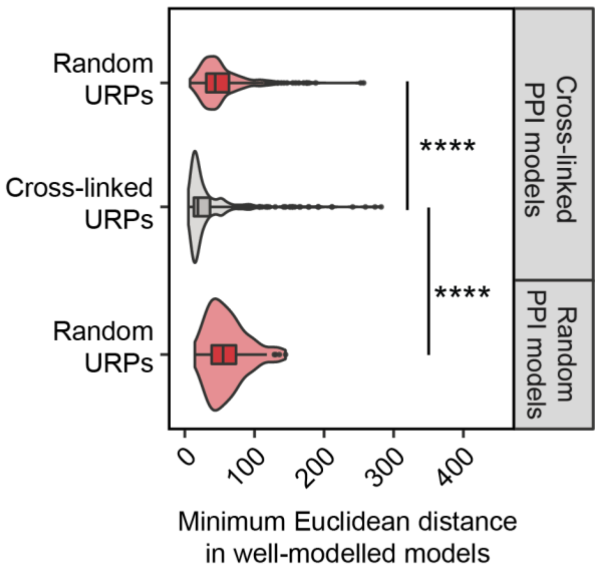


**Supp Figure 9. AlphaFold Multimer-v2 modelling crosslinking statistics. (A)** The distribution of measured Euclidean distances (Å, Cα-Cα) of experimentally derived and randomly generated URPs. Only URPs involving high-confidence residues with pLDDT ≥ 70 were used for model assessment. Experimentally derived URPs were found to be significantly shorter than randomly generated URPs. . **** is $p = 2.2 \times 10^{-16}$.

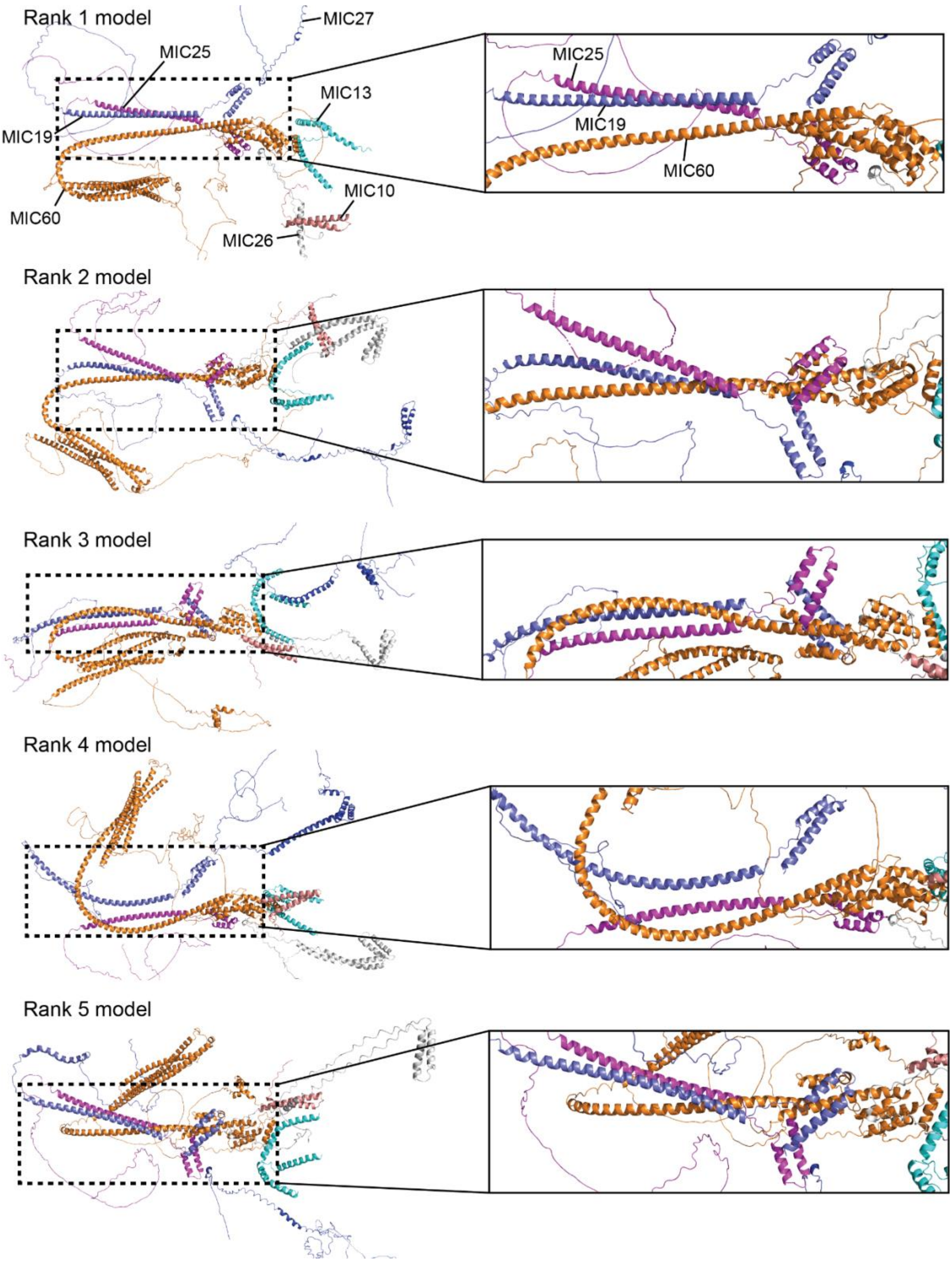


**Supp Figure 10. AlphaFold Multimer-v2 models of the MICOS complex have generally similar core structures comprised of MIC60, MIC25 and MIC19.** While the five ranked AF Multimer-v2 models do not have an overall consensus structure for the heptameric complex, the spatial arrangement of MIC60 (*orange*), MIC25 (*magenta*) and MIC19 (*slate*) were generally similar. Enlarged images of the MIC60-MIC25-MIC19 core (*dashed boxes*) are shown on the right of each model.
